# Spatial organization of the mouse retina at single cell resolution

**DOI:** 10.1101/2022.12.04.518972

**Authors:** Jongsu Choi, Jin Li, Salma Ferdous, Qingnan Liang, Jeffrey R. Moffitt, Rui Chen

## Abstract

The visual signal processing in the retina requires the precise organization of diverse neuronal types working in concert. We performed spatial transcriptomic profiling of over 100,000 cells from the mouse retina, uncovering the spatial distribution of all major retina cell types with over 100 cell subtypes. Our data revealed that the retina is organized in a laminar structure at the major cell type and subgroup level, both of which has strong correlation with the birth order of the cell. In contrast, overall random dispersion of cells within sub-laminar layers indicates that retinal mosaics are driven by dendritic field patterning rather than neuron soma placement. Through the integration of single cell transcriptomic and spatial data, we have generated the first comprehensive spatial single cell reference atlas of the mouse retina, a resource to the community and an essential step toward gaining a comprehensive understanding of the mechanism of retinal function.

**Graphical Abstract:** 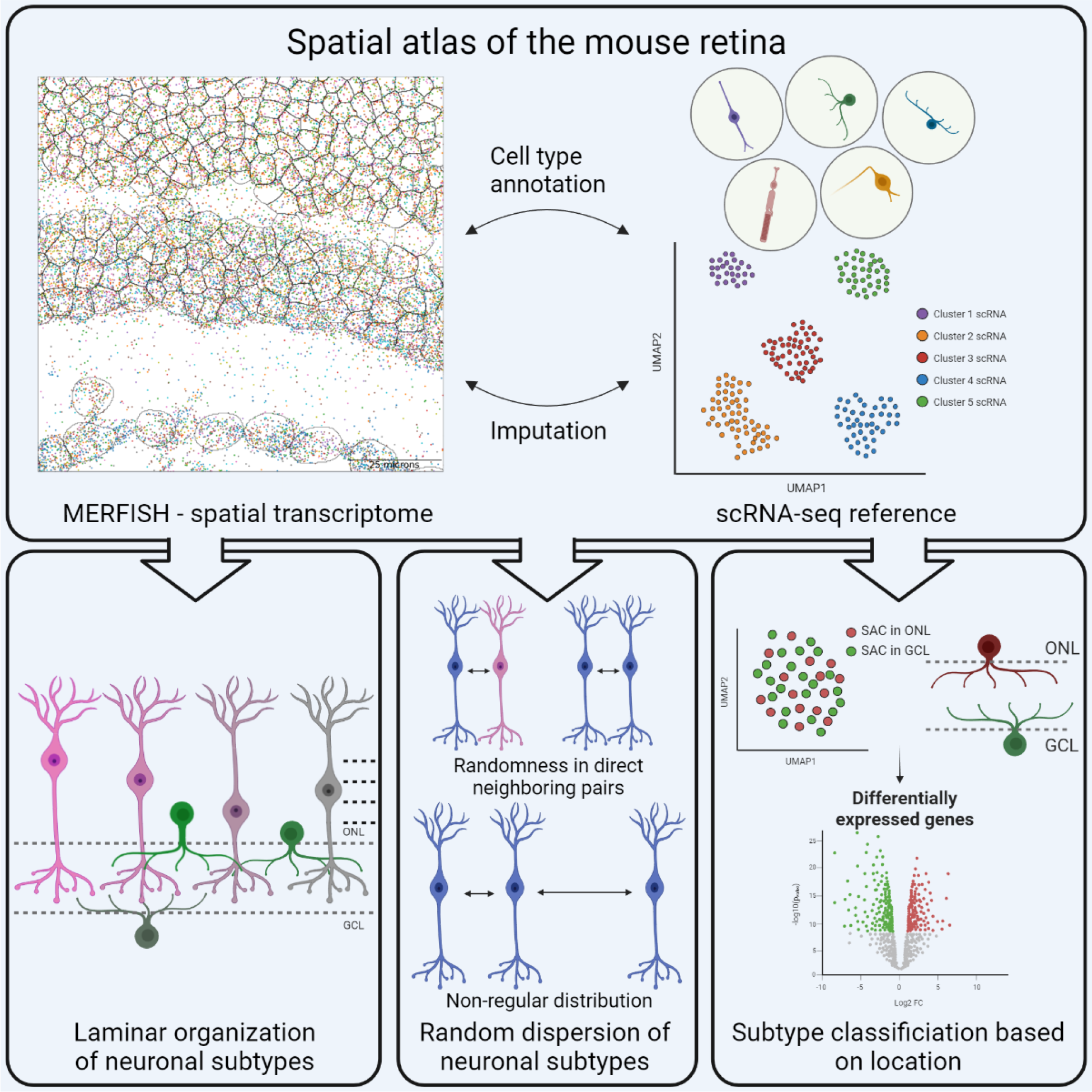

## Introduction

Highly patterned organization of cell bodies and synaptic connections among neurons is essential for the proper function of the central nervous system (CNS). As a part of the CNS, the retina tissue captures and processes the light signal before relaying to the visual cortex. This complex process is achieved through the coordinated action among diverse types of neurons, each with distinct characteristics. The retina is composed of five major neuronal cell types, which can be further classified into many subtypes based on their distinct morphology, gene expression, and function. Within the mouse retina, 128 distinct neuronal subtypes have been identified based on single cell transcriptomics profiling (Shekhar et al., 2016; Tran et al., 2019; Yan et al., 2020). While cataloguing the large number of neuronal subtypes is important in understanding the retina function, charting the organization and interaction among these neurons are equally critical.

Thorough investigation of the synaptic connections in plexiform layers has revealed stereotypical patterns of dendrites and axons of each retinal cell types that target specific sub- layers of the plexiform layer (Amini et al., 2018; West and Cepko, 2022). In contrast, the spatial organization of the neuronal nuclei is less well characterized. On the major cell type level, the laminar organization of nuclei is evident. The distribution of the nuclei on the cell subtype level, on the other hand, has not been systematically investigated. Previous studies indicate that nuclei of some cell subtypes may also follow a general laminar organization. For example, the two amacrine cell (AC) subgroups, GABAergic and glycinergic ACs, show distinct sublayer localization within the inner nuclear layer (INL) with GABAergic ACs basally positioned closer to the ganglion cell layer (GCL) in general (Voinescu et al., 2009). Furthermore, it has been reported that the laminar position of the cell nuclei could affect its function and neurites remodeling of interacting cells (Burger et al., 2021; Nemitz et al., 2021; Sonntag et al., 2012).

Although recent progress in single cell transcriptomic technologies have generated a near complete molecular map for all cell subtypes in the retina, the current droplet-based methods require the dissociation of the tissue into single cells prior to profiling, resulting in the loss of spatial information (Chen et al., 2021; Macosko et al., 2015). Thus, how the 128 retinal cell subtypes are spatially organized in the mouse retina is not well known. Given the interplay between the location and the function of the cell, having a matching spatial map of the retina at the single cell level is critical. So far, the spatial information of cell subtypes is primarily generated based on antibody staining and in situ hybridization of cell type specific markers. Due to the large number of cell types, coupled with the fact that most cell subtypes do not have specifically unique markers, these traditional approaches are not scalable or feasible to generate a complete spatial map. Over the last decade, rapid progress has been made in developing single cell spatial transcriptomic technologies and analyses (Cho et al., 2021; Dries et al., 2021a), including the multiplexed error-robust fluorescence in situ hybridization (MERFISH) technology that can profile 100s to 1000s transcripts directly on tissue sections at single cell resolution (Chen et al., 2015; Moffitt et al., 2018; Rao et al., 2021; Zhuang, 2021). Several recent studies have applied MERFISH to generate single cell spatial maps for various tissues, such as the brain and liver (Lu et al., 2021; Zhang et al., 2021).

In this study, we aimed to generate a spatial atlas of the mouse retina at single cell resolution. Using MERFISH, we profiled over 100,000 single cells from four adult wild type mice retina using a panel of 368 marker genes, which allowed identification of over 100 cell subtypes. The spatial distribution of identified cells revealed that cells from the same subtypes tend to occupy the similar sublayer, and the laminar layer positioning of cell subtypes has a strong correlation with their birth order, suggesting a connection between the laminar structure formation and developmental timing. Within specific laminar layers, we observed random dispersion of cell at the subtype level, indicating that most neuronal subtypes are not assembled in regular arrays and that the retinal mosaics are driven by dendritic field patterning instead of neuronal soma placement. Lastly, the integrative analysis of MERFISH and single cell RNA sequencing (scRNA-seq) data allowed imputation of the entire transcriptome across the retina. Interestingly, we observed differential gene expression among displaced AC subtypes located between INL and GCL, highlighting the importance of spatial information in cell type classification. In summary, the first spatially resolved single cell reference map of the retina generated in this study serves as a valuable resource for the entire vision science community and lays the foundation for many future studies, including development, cell-cell interaction, and circuitry.

## Results

### Single cell spatial transcriptome profiling of the mouse retina

The precise organization of neuronal soma, which presumably affects synaptic connections, is critical for working retina circuitry. To investigate the intricate spatial relationship of retinal neurons on the single cell resolution, we performed spatial transcriptome profiling of the mouse retina tissue using MERFISH (Figure 1A). Based on published scRNA-seq data, a panel of 368 transcripts specific for major cell types and cell subtypes was selected to capture the cell heterogeneity in the mouse retina (Figure 1B and supplemental table 1).

**Figure 1.**
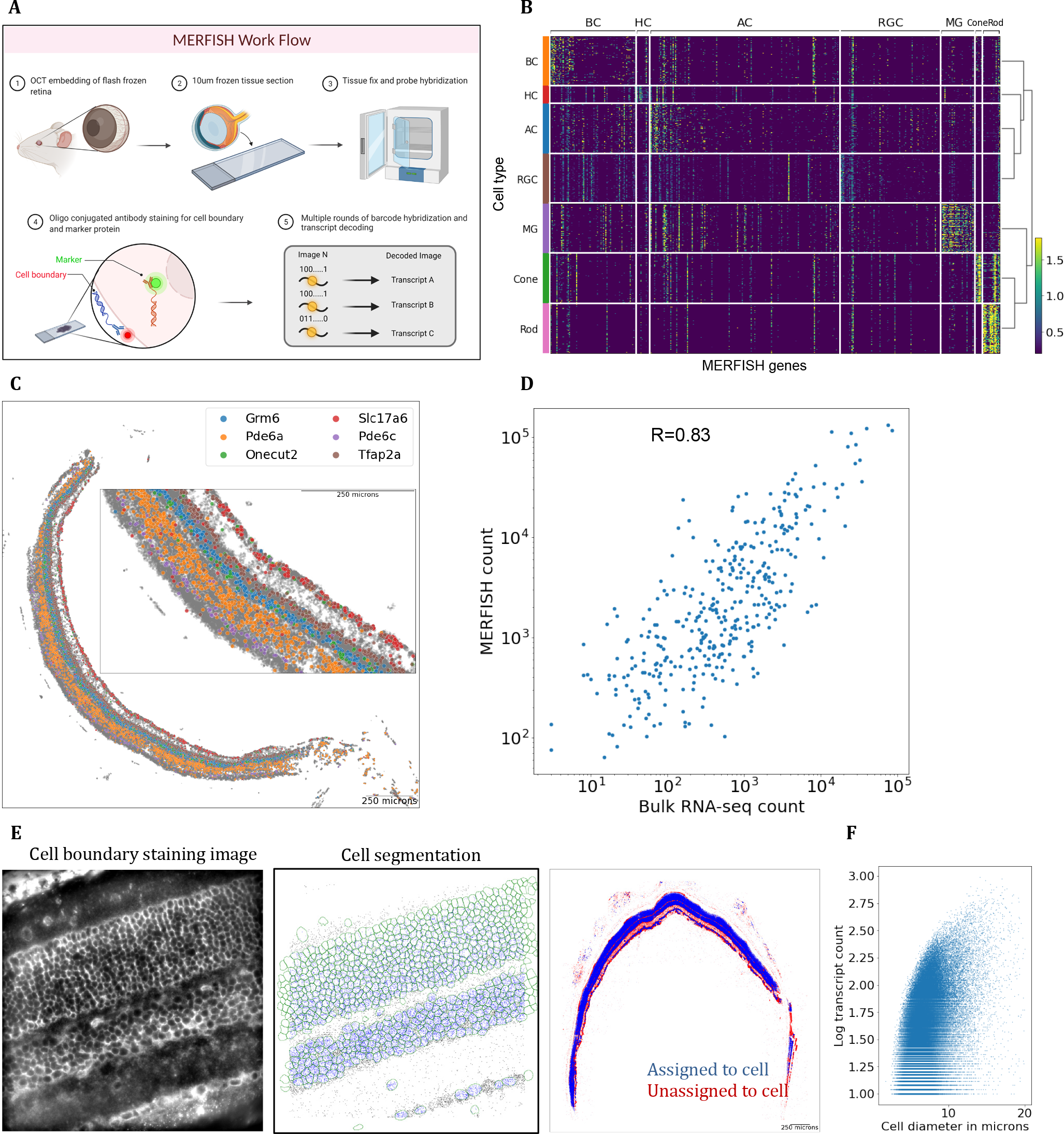
Overview of spatial transcriptome profiling in the mouse retina. **(A)** Overview of the MERFISH protocol workflow. MERFISH experiment was conducted on 10-µm-thick mouse retina cross-sections. Cell membrane staining and in-situ hybridization of transcripts were imaged through multiple rounds readout hybridization. **(B)** Heatmap of MERFISH probe transcript expression in reference scRNA-seq data. Total of 368 markers specific for major cell types and subtypes was selected from scRNA-seq data. **(C)** Distribution of major cell type markers in the retina tissue. Decoded transcripts show expected patterns across the tissue such as photoreceptor markers Pde6a and Pde6c in the ONL, interneuron markers Grm6, Tfap2a, and Onecut2 in the INL, and ganglion cell marker Slc17a6 in the GCL. **(D)** Transcript count comparison between MERFISH and bulk RNA-seq. A high correlation shows MERFISH transcription detection level is comparable to RNA-seq. **(E)** Cell boundary staining image and segmentation result. **(F)** Scatter plot of cell diameter and transcript counts per each MERFISH cells. The single-cell QC metrics shows ∼80 transcript counts per cell with ∼7-µm diameter on average.

Following the workflow in Figure 1A, we performed MERFISH experiments on a total of 28 retinal cross-sections. A high correlation coefficient of 0.98 between tissue sections and experimental replicates indicated excellent reproducibility (Figure S1A). To further evaluate the data quality, we examined the spatial distribution of several known cell type marker genes, which showed proper localization such as photoreceptor markers Pdec6a and Pdec6c in the outer nuclear layer (ONL), bipolar cell (BC) and AC markers Grm6 and Tfap2a in the INL and retinal ganglion cell (RGC) marker Slc17a6 in the GCL (Figure 1C). Furthermore, the transcript count obtained in our MERFISH experiments and the bulk RNA-seq showed positive correlation with a large Pearson correlation coefficient of 0.83, indicating that the overall transcript level measured by MERFISH and RNA-seq are consistent with one another (Figure 1D).

Proper segmentation of cells is challenging as the retina tissue is densely packed with small soma, and nuclei staining alone is not sufficient to identify individual cells. By including the cell membrane co-staining in MERFISH experiments, we employed deep-learning segmentation algorithms to determine cell boundaries and identify individual cells (Greenwald et al., 2022; Stringer et al., 2021) (Figure 1E). Upon segmentation, a total of ∼110,000 cells were obtained (Figure 1E). The average diameter of identified cells was ∼7µm, which corresponds well with other immunohistochemistry images (Figure 1F). The mean number of assigned transcripts per cell was around 80 (Figure 1F).

Integrative clustering analysis (Lopez et al., 2018) of the MERFISH experiments identified 21 retinal and 4 non-retinal clusters (Figure 2A, Figure S1B and C). The retinal single-cell clusters were annotated as one of the six major cell types based on marker expression such as Pdec6a for rods, Pdec6c for cones, Vsx2 for BCs, Pax6 for ACs, Vim for Muller glial cells (MGs), Onecut1 for horizontal cells (HCs), and Slc17a6 for RGCs (Figure 2B). The non-retinal clusters were labeled as RPE, immune cells, astrocytes, and microglial cells based on cell coordinate and gene (Figure 2A, Figure S1C). The proportion of major cell types was consistent with the known composition of mouse retina with rod cells comprising almost 60% of the total population and other INL cell types, cone, and RGC subsequently trailing (Figure 2C). The cell type annotation was confirmed by back-plotting the cell coordinates, which demonstrates consistent layering patterns of major cell types (Figure 2D).

**Figure 2.**
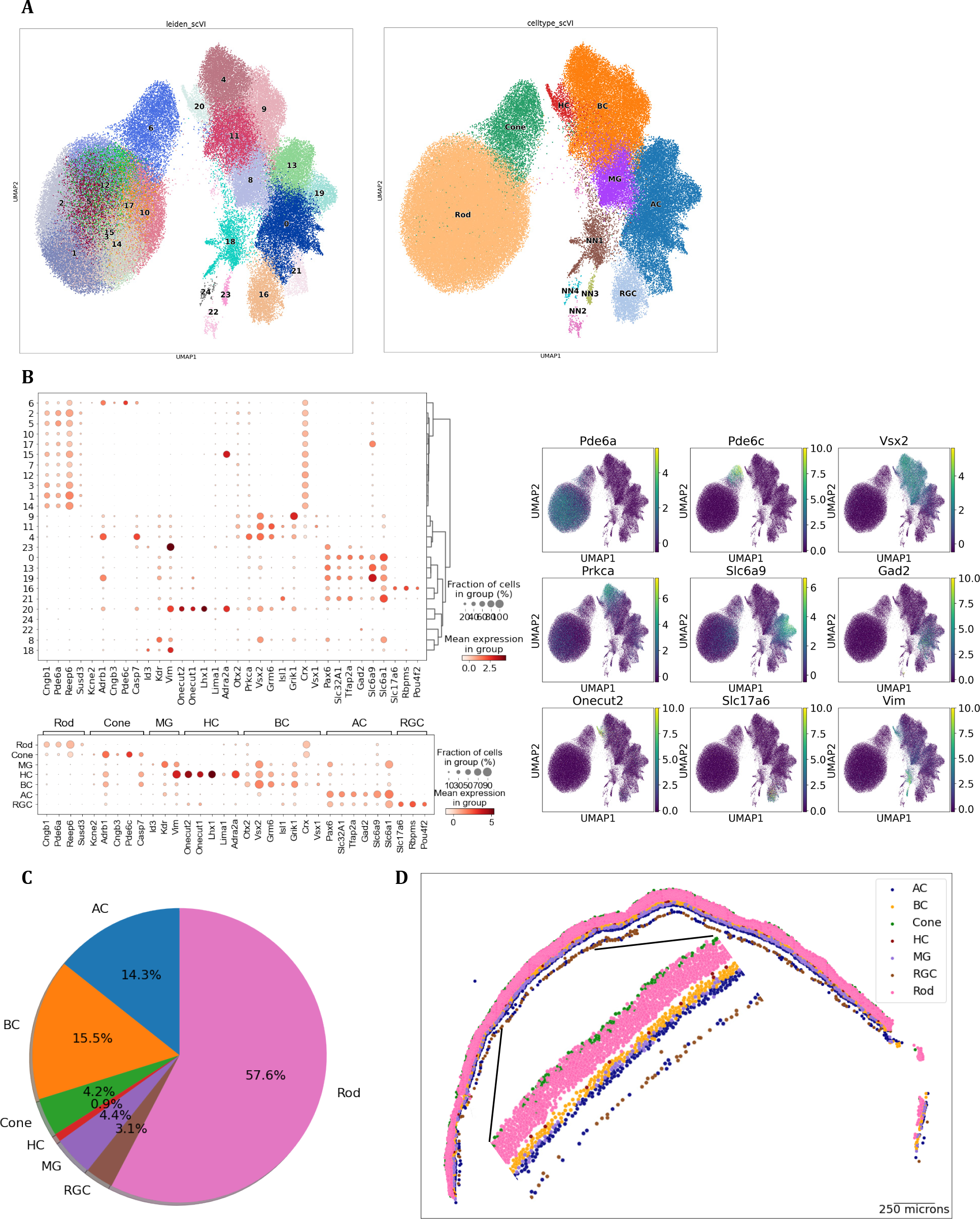
Major cell type identification of MERFISH single-cell profiles. (A) **Visualization of** integrated MERFISH single-cell clusters by UMAP. Integrated clustering analysis of four MERFISH experiments resulted in 21 distinct retinal clusters, which have been annotated as major retinal cell types and 4 clusters, which have been annotated as non-major retinal cell types. **(B)** Major cell type marker expression in retinal single-cell clusters. Cell type specific markers show distinct and exclusive expression in each cluster. Following transcripts were used to annotated cell type: Rod - Cngb1, Pde6a, Reep6, and Susd3; Cone - Kcne2, Adrb1, Cngb3, and Pde6c; BC - Otx2, Vsx2, Isl1, and Grik1; AC - Pax6, Slc32a1, Tfap2a, Gad2, and Slc6a9; RGC - Slc17a6, Rbpms, and Pou4f2; HC - Onecut1, Onecut2, and Lhx1; MG - Id3, Kdr, and Vim. **(C)** Composition of major cell type annotation. The ratio of major cell types matches known cell type composition. **(D)** Tissue plot showing annotated major cell type. Major cell types can be found in appropriate retinal layers.

### Spatial map of over 100 retinal cell subtypes is identified through integration with scRNA-seq data

Five major neuronal cell types in the retina can be further classified into over 100 distinct cell subtypes with unique molecular characteristics. The 15 distinct BC subtypes in the mouse retina can be grouped in three major groups, ON rod, OFF cone and ON cone types based on their functional circuitry (Shekhar et al., 2016). Consistently, in our dataset, the ∼20,000 BCs could be separated into three clusters corresponding to rod (Prkca+), ON cone (Grm6+), and OFF cone types (Grik1+) (Figure S2A). Despite the 3-4 markers per each BC subtype included in the panel, the 368 genes were not sufficient to clearly cluster and separate all subtypes. To achieve better resolution, we leveraged scRNA-seq data to perform co-embedding and integration (Dou et al., 2022) and identified all 15 subtypes with high resolution and minimal batch effect (Figure 3A, Figure S2B, Figure S2C, see Methods and Resources for details). Each cluster of annotated MERFISH BC subtypes exhibited clear and exclusive expression of known subtype markers (Figure 3B). In addition, the resulting cell subtype proportion was similar to the previous estimate based on the scRNA-seq data (Figure 3C) (Shekhar et al., 2016).

**Figure 3.**
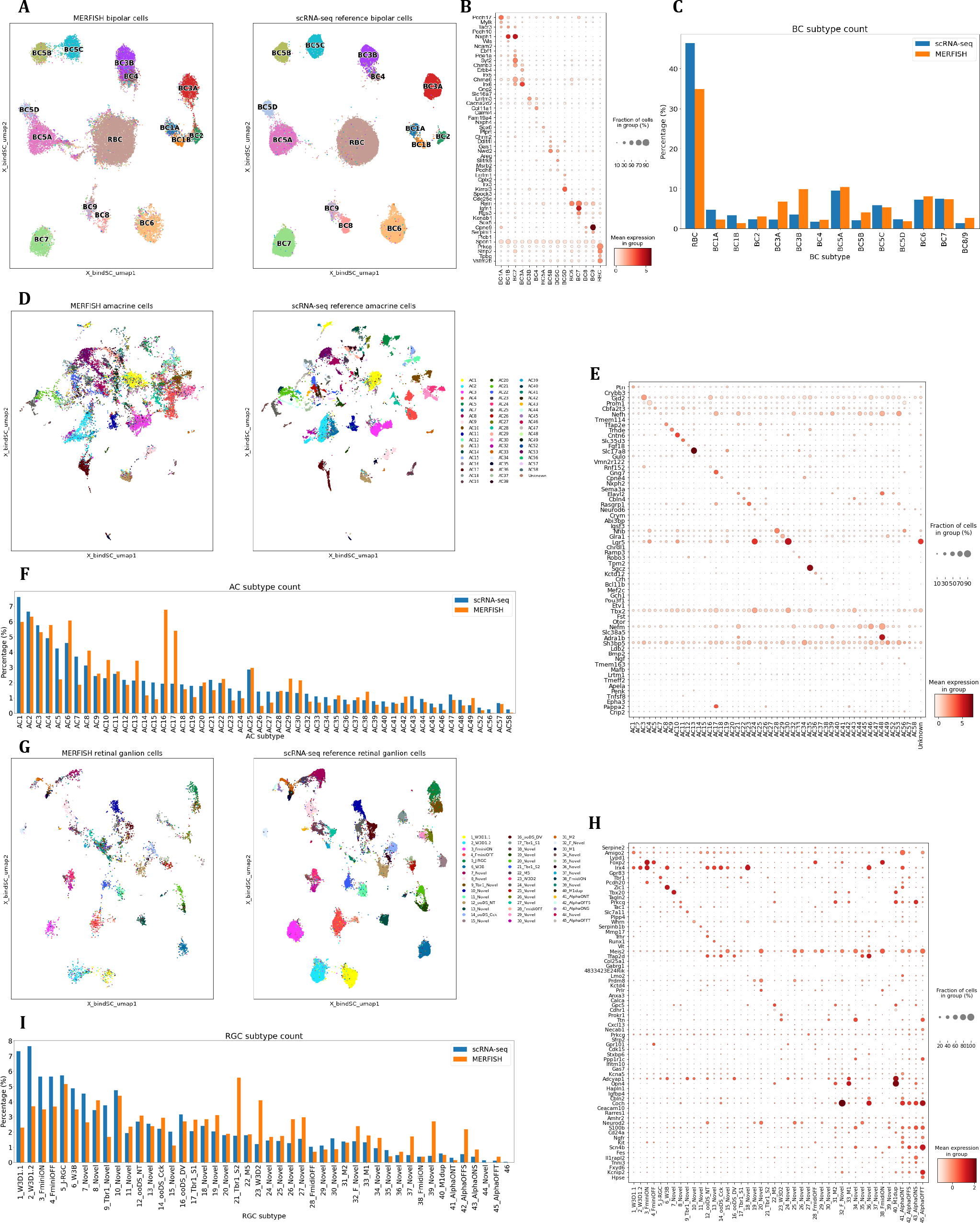
Subtype identification of MERFISH cells by integration with scRNA-seq reference. **(A)** Co-embedding plot of bipolar cells between MERFISH and scRNA-seq. A high-resolution clustering map is obtained by integration between MERFISH (left) and scRNA-seq (right) with strong overlap. **(B)** Dot plot of bipolar cell subtype marker expression in annotated MERFISH subtypes. **(C)** Bar plot of annotated bipolar cell subtype ratio. The ratio of annotated BC subtypes show similarity with published scRNA-seq study. **(D)** Co-embedding plot of amacrine cells. Integration between MERFISH (left) and scRNA-seq (right) ACs show reasonable overlap and separation from each subtype cluster. **(E)** Dot plot of AC subtype marker expression. **(F)** Bar plot of annotated AC subtypes ratio. The ratio of annotated AC subtype is largely consistent with published scRNA-seq study with a couple of exceptions such as increased starburst amacrine cells in the MERFISH experiment. **(G)** Co-embedding plot of retinal ganglion cells. Integration between MERFISH (left) and scRNA-seq (right) RGCs shows strong overlap. **(H)** Dot plot of RGC subtype marker expression. **(I)** Bar plot of annotated RGC subtype ratio. The ratio of annotated RGC is comparable with the published scRNA-seq study.

A total of ∼17,000 cells were annotated as AC by the expression of pan-AC markers Pax6, Tfap1 and Slc32a1 (Figure 3D). Following the initial pan-AC assignment, we first accessed the canonical GABA and glycine neurotransmitter markers Slc6a1 and Slc6a9. Similar to BC, confined expression of Slc6a1 and Slc6a9 was observed across the substructures of 4 AC clusters (Figure S2D). Through the co-embedding analysis, we achieved a significantly higher resolution map, allowing the annotation of MERFISH AC clusters to its corresponding subtype (Figure 3D, Figure S2E, see Methods and Resources for details). Consistently, MERFISH AC subtypes showed clear expression patterns of known subtype markers (Figure 3E) (Yan et al., 2020). Moreover, overlaying the subtype annotation from the co-embedding result on the lower dimensional space calculated using only MERFISH features showed distinct and confined sub- structures of subtype labels, providing further confidence to our prediction (Figure S2F). The population abundance of AC subtypes identified ranged from 0.03-6.78% with a decreasing trend to the rarer subtype, largely consistent with the estimates based on scRNA-seq data (Yan et al., 2020) with a few exception such as increased ratios of AC16 and AC17 (Figure 3F).

The expression of RGC markers such as Slc17a67 and Rbpms were used to annotate RGCs in our data (Figure 3G). The total number of annotated RGC was ∼3,400 and comprised 3.1% of the total population (Figure 2C). Upon co-embedding with scRNA-seq reference data, 44 out of 46 RGC subtypes were annotated (Figure 3G) (Tran et al., 2019). Examination of the known combinatory RGC subtype markers (Tran et al., 2019) indicated that our RGC subtype annotation shows comparable subtype specific marker expression (Figure 3H). The proportion of annotated RGC subtypes in our dataset was reasonably correlated with previous estimate (Figure 3I) (Tran et al., 2019). As expected, annotated RGCs showed expected localization exclusively in the GCL (Figure S2G).

### Neuronal subtypes show distinct laminar positions

Laminar organization of major cell type soma is critical for proper the visual circuitry and function of the retina; however, it is not as well understood how soma are organized on the subtype level. To investigate the positioning patterns of neuronal subtypes, we calculated cell position along the normalized INL length. As shown in Figure 4A, on average, rod ON bipolar cells (RBCs) exhibited the most apical cell body relative to other BC subtypes (p-value 0.001), positioned within the top 20% of the INL length. In addition, RBCs showed a strong proximity enrichment with HCs, the most apical cell type in the INL (Figure S3A). This observation agrees with the previous reports that show RBC immunostaining closest to the outer plexiform layer (OPL) (Greferath et al., 1990; Haverkamp et al., 2003). In contrast, BC1B, a rare population of OFF BC subtype (Della Santina et al., 2016; Shekhar et al., 2016), showed the largest distance away from the OPL relative to other BC subtypes (Figure 3D, Figure S3B, p value 0.001).

**Figure 4.**
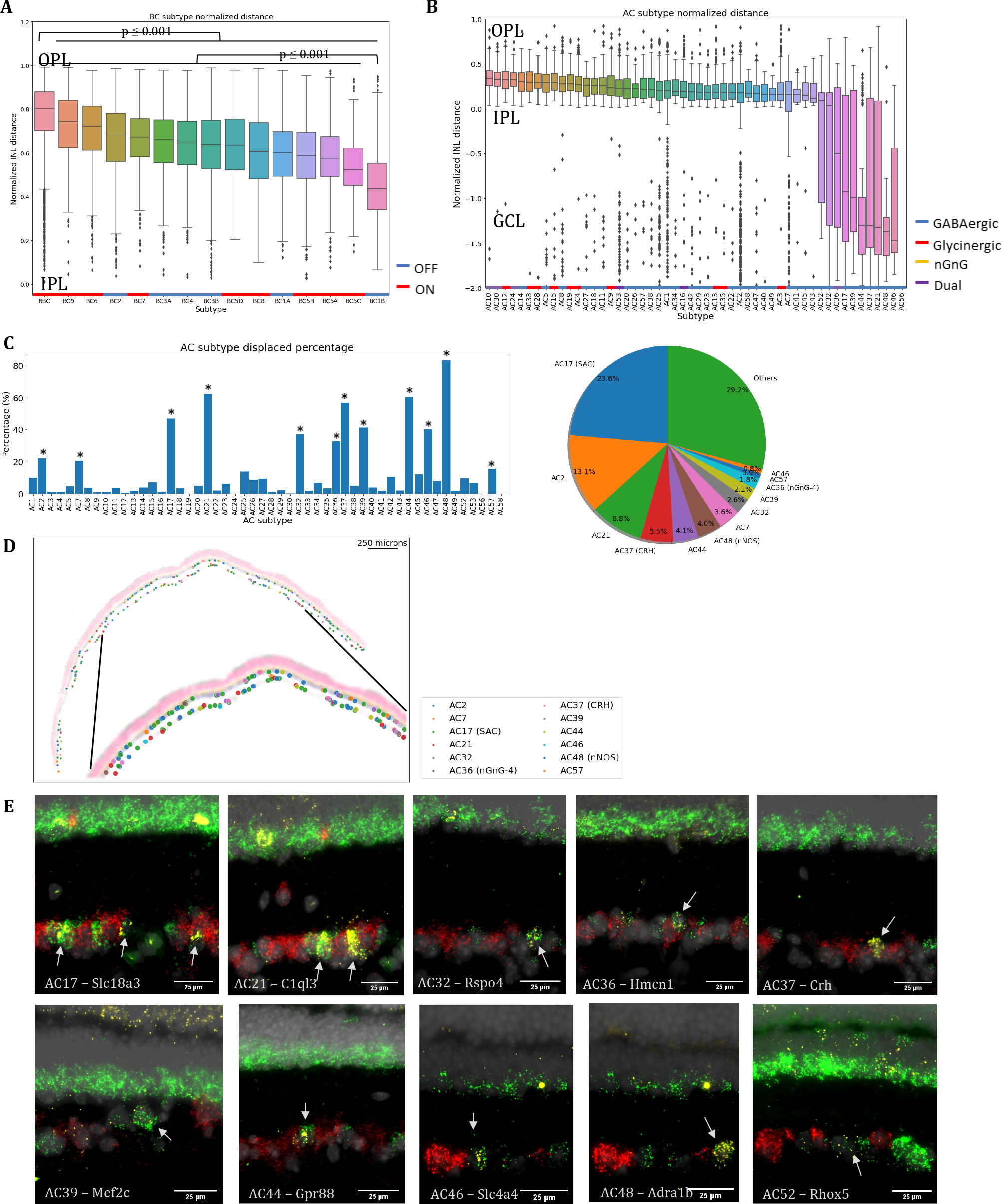
Laminar organization of neuronal subtypes in the retina. **(A)** Boxplot of bipolar subtype position in the normalized INL length. BC subtypes showed overlapping, yet distinct positioning patterns (ON types in red, and OFF types in blue). RBCs were positioned most apically compared with other subtypes (p value < 0.001), whereas BC1B showed significant basal positioning against other subtypes (p value < 0.001). OFF and ON BC subtypes are marked with blue and red bars, respectively **(B)** Boxplot of amacrine subtype position in the normalized INL length. Most AC subtypes showed distinct positioning within the bottom half of the INL (GABAergic in blue, glycinergic in red, nGnG in yellow, and dual in purple). GABAergic, glycinergic, nGnG, and dual subtypes are marked with blue, red, yellow, purple bars, respectively. **(C)** Bar plot of displacement proportion in each amacrine subtype and a pie chart of displaced subtype composition in the GCL. Twelve AC subtypes showed significant displacement (p-value < 0.01) and made up more than 70% of all ACs in the GCL. **(D)** Tissue plot of displaced amacrine subtypes. Displaced AC subtypes showed distribution across both INL and GCL. **(E)** In-situ hybridization images of displaced AC subtype markers in the ganglion cell layer. Specific markers against ten displaced AC subtypes were profiled in conjunction with pan-AC marker Slc32a1 and pan-RGC marker Slc17a6. Specific subtype markers in yellow showed co-localization with the pan-AC marker in green, but not with the pan-RGC marker in red.

Located toward the central INL, BC1B showed a high proximity enrichment with MGs (Figure S3B). As BC1B has been shown to possess amacrine-likeness in morphology and positions (Della Santina et al., 2016; Shekhar et al., 2016), the preferential BC1B positioning in the center of the INL is in agreement with literature. Interestingly, although no difference between the average ON and OFF cone bipolar cell position was found (Figure S3C), each BC subtype showed overlapping yet distinct sublaminar distribution (Figure 4A). Specifically, certain subtypes such as BC9 and BC6 exhibited more apical location, while subtypes like BC5C and BC5A showed more basal positioning. It has been observed that the birth timing of cone BCs precede RBCs in general (Morrow et al., 2008), and the OFF cone subtypes are born significantly earlier than the ON cone subtypes (West et al., 2022). The general trend of apically positioned RBC and a few ON subtypes and a few basally positioned OFF subtypes suggests that the laminar organization is correlated with but not entirely determined by the birth order of BC subtypes.

ACs can be grouped into four larger subgroups, GABAergic, glycinergic, non-GABAergic and non-glycinergic (nGnG) (Kay et al., 2011), and dual receptors (Yan et al., 2020). Previous studies suggest that GABAergic ACs are the earliest born types and their cell bodies are located at a more basal position relative to glycinergic ACs (Cherry et al., 2009; Voinescu et al., 2009). The comprehensive annotation of AC subtypes in our study revealed that AC subtypes possess overlapping yet distinct position within the INL (Figure 4B). Consistent with previous observations, we observed that glycinergic subtypes were more apically positioned whereas GABAergic subtypes were more basal. Interestingly, three nGnG subtypes (AC10, AC24, and AC30) were among the most apically positioned subtypes. The three nGnG subtypes share transcriptional similarity with glycinergic subtypes and cluster within the glycinergic clade (Yan et al., 2020). Several AC subtypes showed significant cell distribution outside of the normalized INL length, presumably representing displaced amacrine subtypes.

### Twelve amacrine cell subtypes are preferentially displaced in the ganglion cell layer

About half of the GCL is composed of ACs out of their endogenous location in the INL, named displaced ACs (Jeon et al., 1998; Pang and Wu, 2011). Several displaced AC subtypes such as starburst, CRH+, VIP+, nGnG, and nNOS+ subtypes have been identified based on their morphologies and marker labeling (Akrouh and Kerschensteiner, 2015; Jacoby et al., 2015; Zhu et al., 2014) . However, it remains unknown exactly which subtypes are preferentially displaced with the recent molecular classification of 63 AC subtypes (Yan et al., 2020). To identify preferentially displaced AC subtypes, we determined the proportion of ACs mapped to the GCL and calculated the likelihood of each subtype to be displaced (Figure 4C). Our analysis revealed 12 AC subtypes with significant displacement, total displacement ratio ranging from 23% to 80% (Figure 4C, 4D, p value < 0.01). Consistent with previous observation, the starburst amacrine cell (SAC) was the most abundant displaced AC type and accounted for 24% of all displaced ACs (Figure 4C) (Vaney et al., 2012). Furthermore, CRH1 (AC37), nGnG-4 (AC36), and nNOS (AC48) were also identified to be displaced, confirming the previous report (Akrouh and Kerschensteiner, 2015; Jacoby et al., 2015; Zhu et al., 2014). Out of 12 displaced ACs, 8 subtypes have not been reported as displaced ACs. It is worth noting that with the exception of AC36 (nGnG-4), the remaining displaced subtypes ACs are all GABAergic. To validate the findings, we performed RNA in-situ hybridization against molecular markers specific to the novel displaced AC subtypes in conjunction with pan-AC marker Pax6 and pan-RGC marker Rbpms. As shown in Figure 4E, cells positive for AC subtype markers and Pax6, but negative for Rbpms were observed in both the INL and the RGC layer, validating their identity as displaced ACs.

### Most neuronal subtypes are randomly dispersed in laminar sublayers

The dendrites of retinal neurons are widely believed to be assembled in an ordered pattern that allows the most efficient coverage of the tissue by diverse neuronal types, termed retinal mosaics (Jeon et al., 2007; Kay et al., 2012; Raven and Reese, 2002; Reese and Keeley, 2015, 2016; Wässle et al., 1978). To investigate the local intracellular spacing rules that govern the distribution of retinal neuron soma, we applied two different statistical tests to determine the properties of retinal mosaics on the subtype level. We first performed proximity enrichment analysis (Dries et al., 2021b) to identify the nearest neighbor relationship of neuronal subtypes. To identify nearest neighbor relationships across all cells, we constructed a pairwise cell-cell network in our spatial map (Figure S4) and performed subtype label-shuffling of cells within binned INL lengths to calculate the observed over expected frequency of cell-cell pair occurrence. We observed that the overall cell-cell neighboring pairs were randomly distributed with a few exceptions in homotypic and heterotypic pairs. Within the INL, we observed a significantly enriched homotypic pair of BC9 and several enriched homotypic pairs as well as enriched and depleted heterotypic pairs of amacrine subtypes (Figure 5A and 5B, Figure S5A). In the GCL, we found significant enrichment in 7 RGC homotypic pairs, 3 heterotypic RGC pairs, 3 heterotypic AC pairs, and 1 heterotypic pair between AC17 and RGC 20_novel (Figure S5B). The enrichment and depletion of cell-cell pairs of rare subtypes may suggest preferentially attraction or repulsion mechanism driven from subtype genesis or neuronal migration.

**Figure 5.**
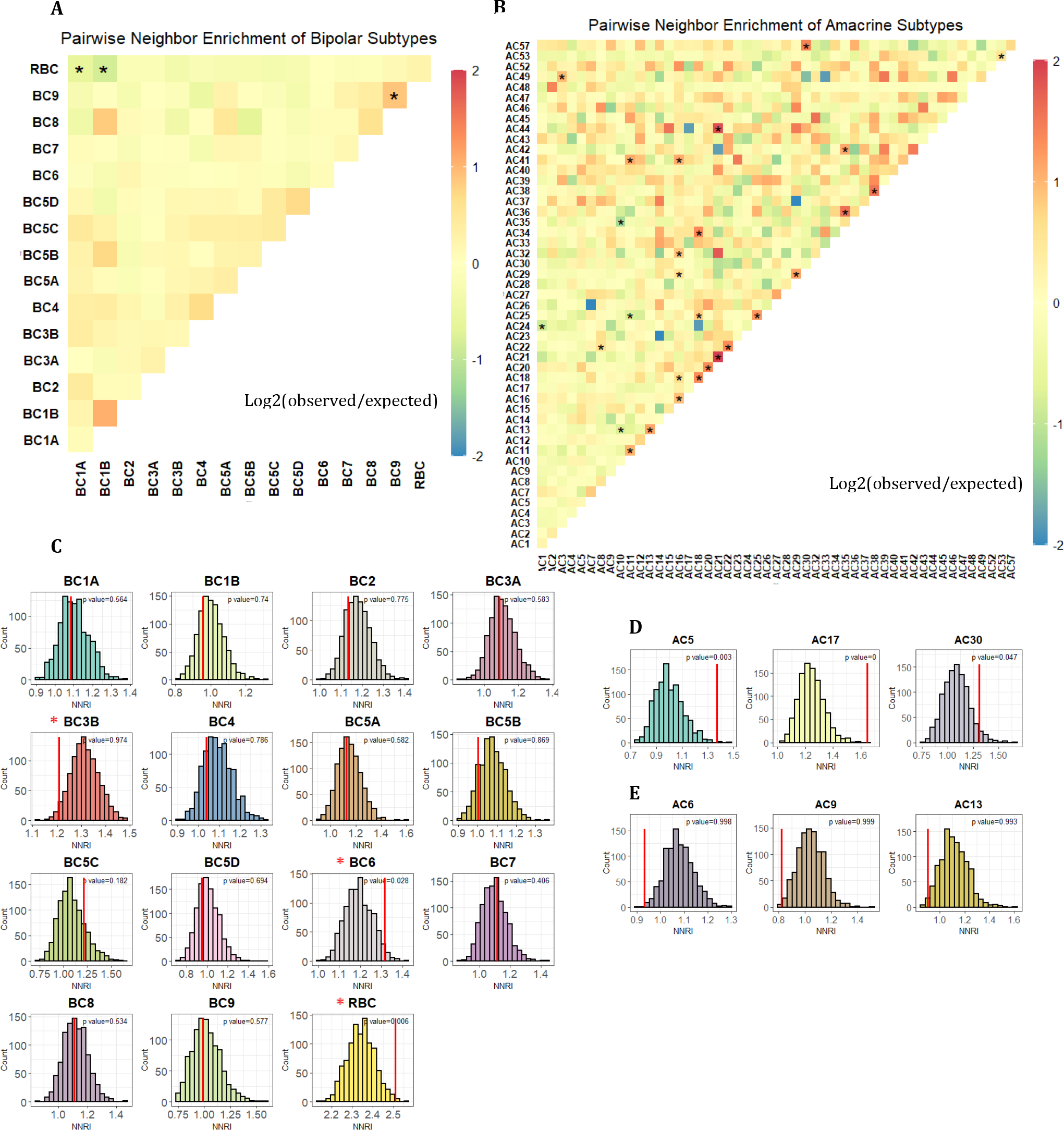
Distribution of neuronal subtypes in the retina. **(A)** Heatmap of pairwise nearest neighbor enrichment scores amongst bipolar subtypes. Statistically significant nearest neighbor pairs are labeled with asterisks. Overall, BC subtypes show random distribution with no enrichment in homotypic or heterotypic subtype pairs. Noteworthy exception is the increased occurrence of homotypic pairs of BC9. **(B)** Heatmap of pairwise nearest neighbor enrichment scores of amacrine subtypes in the INL. Enriched or depleted nearest neighbor pairs with statistical significance are labeled with asterisks. AC subtypes show random distribution with a handful of enriched homotypic pairs and heterotypic pairs. **(C)** Comparison between observed and permuted NNRIs of bipolar subtypes. Observed NNRI of each subtype shown in red line was compared by examining the distribution of permuted NNRIs in each histogram. All but three subtypes exhibited no significant difference in regularity compared with random permutation, suggesting that these neuronal subtypes are randomly distributed. Only BC3B, BC6, and RBC showed significantly decreased or increased regularity indexes and are marked in red asterisks. **(D-E)** Comparison between observed and permuted NNRIs of non-random neuronal subtypes. A few neuronal subtypes exhibited significantly larger **(D)** or smaller **(E)** regularity values compared with random permutation. While larger regularity values show repulsion characteristics, suggesting equal spacing pattern, smaller regularity values show attractive characteristics, suggesting tight cluster formation of cells.

We further calculated nearest neighbor regularity index (NNRI) (Keeley et al., 2020) and performed random label shuffling to examine the regularities in nearest neighbor distance and the self-spacing characteristics of individual neuronal subtypes. The simulation was performed with cells confined within binned INL lengths to compensate for the laminar organization of subtypes. In contrast to the common presumption that retinal neuronal are arranged in an orderly array, we found that most neuronal subtypes exhibit irregular mosaic patterns with no significant difference in distribution regularities from randomly simulation (Figure 5C, Figure S6, S7).

Specifically, we found all but 3 bipolar cell subtypes and 6 amacrine subtypes to follow random distribution in the INL (Figure 5C, 5D, 5E). Among the 9 exceptions, 5 subtypes showed larger average regularity index than permuted, including RBC, BC6, AC5, AC17 (starburst), and AC30 (nGnG-3), while 4 subtypes have smaller average regularity index to the same subtype, including BC3B, AC6 (A17), AC9, and AC13 (VG3) (Figure 5D and 5E p-value < 0.05). Our observation of cholinergic starburst amacrine cells (AC17) with larger NNRI is consistent with previous reports of regular mosaic organization in cholinergic amacrine cells(Galli-Resta et al., 2000; Whitney et al., 2008).

### Spatial map of whole transcriptome can be obtained through imputation

Although the 368 genes were sufficient in annotating cell types and subtypes, spatial expression profile of genes outside of the panel remains unknown. To obtain the spatial information for the entire transcriptome, we performed imputation by leveraging scRNA-seq data and constructed UMAP visualization of cell clusters (Figure 6A, see Methods and Resources for details). In brief, upon co-embedding of MERFISH with scRNA-seq data, imputed gene expression of each MERFISH cell was calculated from weighed expression values of its neighboring scRNA-seq cells in the low dimension space (Biancalani et al., 2021). As an evaluation of the imputation, we examined several major cell type markers in raw MERFISH and imputed transcriptomes and observed appropriate expression patterns in cell clusters and tissue location of cells (Figure 6B, Figure S8A). Furthermore, known markers not included in the MERFISH panel such as Rho for rods, Crx for rods, cones, and BCs, Apoe for MGs, and Scgn in certain BC subtypes also showed proper tissue location (Figure 6C). A high positive correlation was observed between MERFISH and imputed gene expression across four MERFISH experiments (Figure 6D-E, 0.6 PCC). Taken together, we generated a highly accurate spatial transcriptomic map through imputation by integrating scRNA-seq with MERFISH.

**Figure 6.**
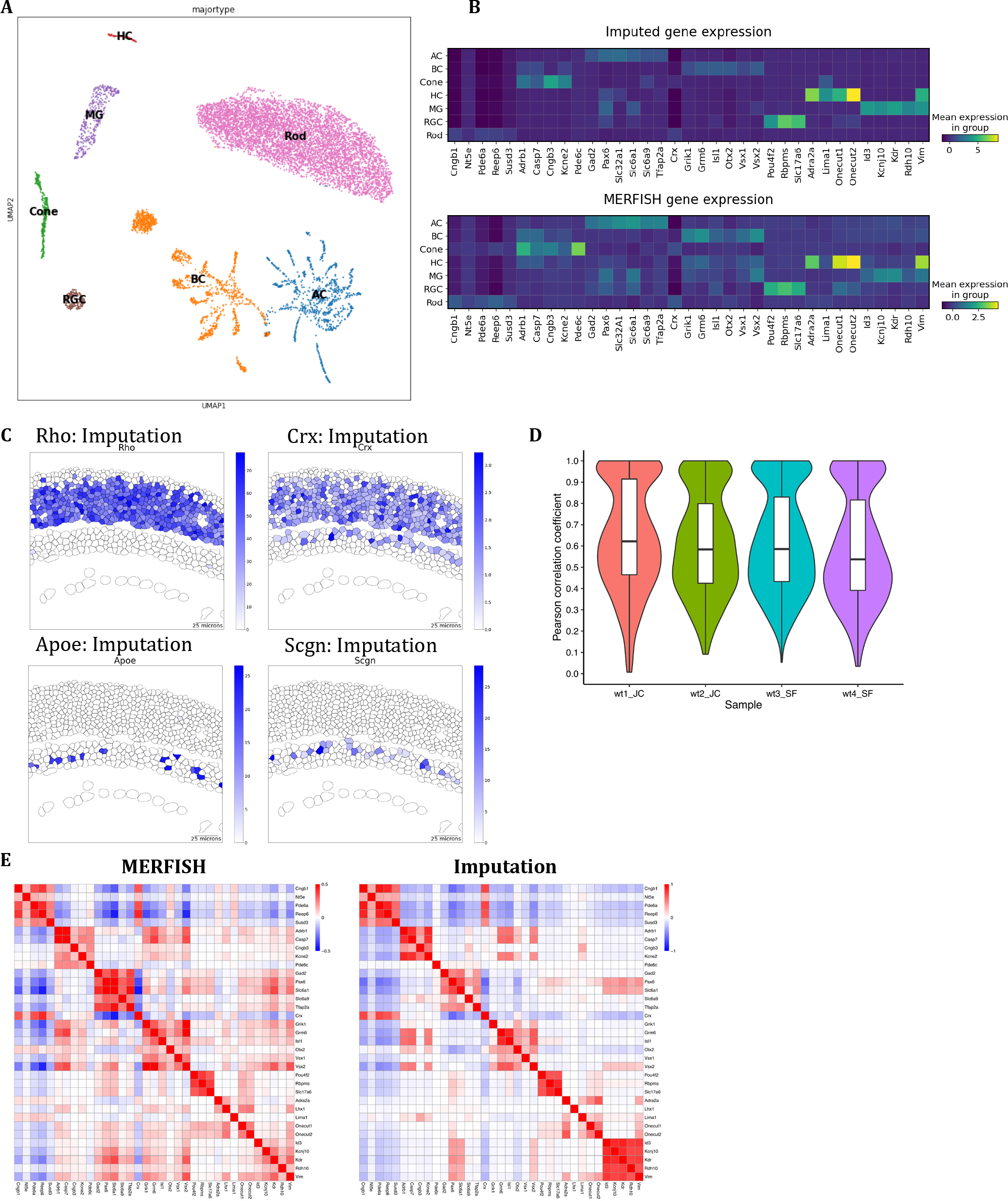
Imputation of the entire transcriptome through integration between MERFISH and scRNA-seq. **(A)** Visualization of single-cell clusters constructed from imputed gene expression. **(B)** Gene expression patterns of major cell type markers in MERFISH and imputation. Both raw MERFISH and imputed transcriptome show proper expression of cell type markers in corresponding cell type clusters. **(C)** Inferred spatial gene expression patterns of cell type markers from imputation. Cell type specific markers such as Rho, Crx, Apoe, and Scgn imputed from scRNA-seq data show appropriate localization. **(D)** Violin plot of correlation coefficient between MERFISH and scRNA-seq by dataset. Positive correlation in all four dataset indicates reliable imputation (PCC 0.6). **(E)** Correspondence between cell type marker expression between MERFISH and imputed transcriptomes. Pearson correlation coefficients between gene expression across cells show close similarities between MERFISH profiles and imputation.

### Gene expression is influenced by the location of the cell

With the entire transcriptome across the retina, we assessed whether the location of the cell affects gene expression within the same cell subtype. We first examined Starburst amacrine cells (SAC) which can be classified into ON and OFF types based on their distinct functions and locations in the GCL and INL, respectively (Famiglietti, 1983). While a distinct transcriptional profile is observed between ON and OFF SACs during maturation, the transcriptional difference in SACs diminishes with age and merge into one scRNA-seq cluster by P18 (Peng et al., 2020).

Consistently, SACs found in the INL and GCL in our experiment were also mapped to one cluster based on the imputed transcriptome (Figure 7A). When differentially expressed gene (DEG) analysis was performed between SACs in found different layers, surprisingly, many genes previously known to be expressed specifically in ON and OFF SACs were identified (Figure 7B, Supplemental table 2). For example, Fezf1, Zfhx4, Slit2, and Cntn5, which are reported to be key transcription regulator or modulate the homophilic interaction with certain RGCs, were enriched in SACs located in the GCL (Figure 7B) (Peng et al., 2020). Within SACs in the INL, genes such as Rnd3, Zfhx3, and Tenm3 were enriched, which support proper OFF SAC function through neurite modulation (Figure 7B) (Peng et al., 2020). In short, our results indicate that distinct transcriptional regulation exists within the same subtype depending on the location of the cell. Our result demonstrates that the spatial information can be used to further divide subtypes into distinct groups with functional and transcriptional differences, which cannot be achieved through scRNA-seq alone.

**Figure 7.**
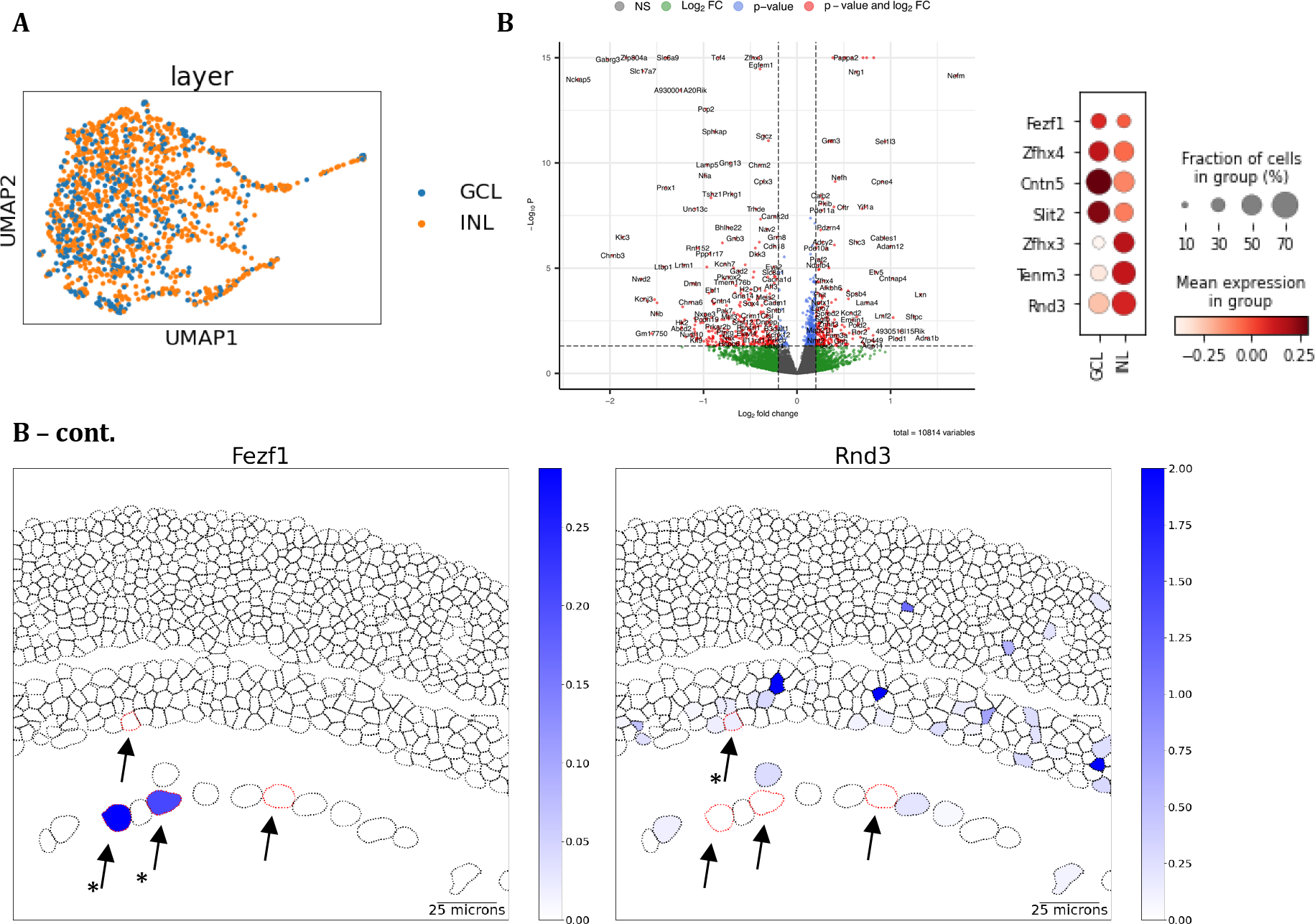
Differential gene expression by location. **(A)** Visualization of merged starburst amacrine cell cluster. Imputed SACs in the INL (orange) and the GCL (blue) are indistinguishable in one cluster. **(B)** Volcano plot of differentially expressed genes between INL and GCL starburst amacrine cells, dot plot of known location specific genes and representative plot of Fezf1 and Rnd3 gene expression plots. Several known genes expressed by ON SAC such as Fezf1, Cntn5, and Slit2 are enriched in the GCL, and genes involved in OFF SACs such as Zfh3, Tenm3, and Rnd3 are enriched in the INL. Starburst amacrine cells are marked with red outlines and arrows in INL and GCL in the tissue plot

### Interactive visualization of the mouse retina spatial atlas

To facilitate the visualization and access of our mouse retina spatial atlas, we developed a user accessible database using the cellxgene software (http://cellatlas.research.bcm.edu) (Megill et al., 2021). The database can be used to visualize the imputed genes on the representative retina tissues (Figure S9). Moreover, the database interface allows examination of pre-computed metadata including annotated major cell types and subtypes. In conclusion, our database provides a user-friendly interactive exploration of the mouse retina spatial atlas, which can serve as a valuable tool for the vision community.

## Discussion

We have generated the first spatially resolved cell atlas of the mouse retina at single cell resolution. Analysis of over 100 cell types and cell subtypes identified in the atlas reveals two principles of cell organization in the retina, lamination and dispersion. We observed stereotypical laminar patterning not only at the major cell type level, but also at the subgroup and subtype level. Within specific laminar sublayers, we observed general randomness in the dispersion pattern of cells belonging to the same subtype classification, indicating that most neuronal subtypes are not arranged in a regular array. Through integration of MERFISH and scRNA-seq data, we constructed a spatial transcriptome map beyond the limited probe panel. By combining the cell coordinates and transcriptome, we discovered twelve displaced AC subtypes, demonstrating the limitation of transcriptomic profiling alone. Furthermore, we observed differential gene expression between ACs of a same subtype in the INL and GCL, providing intriguing evidence of distinct location and function from cells with nearly indistinguishable transcriptome.

### Building a spatially resolved cell atlas of the mouse retina

To distinguish the large numbers of cell types in the highly heterogenous retina, we designed a panel of 368 cell type specific marker transcripts, including 3-4 markers for each BC subtype and 1-2 markers for each AC and RGC subtype based on scRNA-seq data (Shekhar et al., 2016; Tran et al., 2019; Yan et al., 2020). To capture rare cell types, we performed in depth profiling using this panel to identify over 100,000 cells in the mouse retina, including 63,400 rods, 3,300 cones, 20,300 BCs, 17,300 ACs, 1,100 HCs, 5,300 MGs, and 3,400 RGCs and further classified them into over 100 cell subtypes. Furthermore, we have systematically evaluated the quality of our data. First, transcript expression level measured by MERFISH was highly reproducible and concordant with bulk RNA-seq of the retina (Figure 1D). Second, the expression of known marker genes showed an expected pattern in the retina (Figure 1C). Third, distribution of the annotated cell was consistent with previous knowledge (Figure 2D). For instance, the cone cells were frequently observed in the second or third sublayer from the apical ONL. The accurate cell localization pattern extended beyond major cell types, but also at the level of subtypes. As an example, the most common displaced AC subtypes, SACs, were found between both the GCL and the INL. Taken together, this dataset provides a highly sensitive spatial map for most cell types in the retina with high accuracy.

### Integration of MERFISH and scRNA-seq to produce high resolution spatial transcriptome map of the retina

Although the panel of 368 transcript markers was sufficient in annotating most retinal cell types and subtypes, we sought to examine the spatial information of the entire transcriptome through leveraging scRNA-seq data and performing imputation. Thorough evaluation of our imputed results suggests that it is highly accurate. First, high concordance between imputed and measured expression was observed. Second, manually examining the imputed results for a set of known marker genes confirmed expected results. Third, based on imputed data, we were able to identify and confirm differentially expressed genes in displaced SACs compared to SACs in INL. The ability to identify genes with subtle differences in expression pattern underscores the accuracy and robustness of the imputation results. Therefore, our spatial atlas provides not only the cell type information at single cell resolution, but also transcript information of entire transcriptome.

### Organization of cells in the retina

Like the rest of CNS, the retina is highly organized with millions of cells from over hundred different cell types working in concert to efficiently convert and transmit light stimuli.

The organization of cells in the retina appears to follow two prominent rules, laminar structure and mosaic dispersion.

#### Laminar organization of the retina

The characteristic three-layer structure of the retina is formed via the migration of neurons along the radial axis during development, which is critical to the visual circuitry and function (Amini et al., 2018; West and Cepko, 2022). While major cell type occupies a distinct layer of the retina, similar laminar pattern has also been observed for several cell subgroups and subtypes (Voinescu et al., 2009; West et al., 2022). The compendium of spatial distribution on the subtype level generated in this study permitted us to investigate the spatial organization of near all cell types systemically. As such, clear lamination patterns were also observed for both BC and AC subtypes. In our study, we observed preferential localization of RBC at the apical boundary of the INL, consistent with previous knowledge (Figure S3A). The soma distribution of most OFF BC subtypes showed similar depth within the INL except for BC1B, which is located more basal and directly adjacent to the amacrine cells. Interestingly, BC1B exhibits distinct morphology with no direct connection from photoreceptor cells and shows amacrine-like characteristics (Della Santina et al., 2016; Moffitt et al., 2018). The distribution of the cone ON subtypes, on the other hand, was rather dispersed with BC5A, BC5B, and BC5C exhibiting more basal position on average while other subtypes were located more apically. In addition, transcriptionally similar BC8 and BC9 (Shekhar et al., 2016) showed distinct positioning patterns with BC9 exhibiting more apical localization compared with BC8. Although the axon lamination into the inner plexiform layer (IPL) is highly consistent within each BC subgroup and can be used to distinguish between subgroups, the distribution of soma positioning shows a more diverse pattern.

As the most diverse class of neuronal cells in the retina, studies on individual amacrine subtypes are not as well established as BCs. In fact, 6 out of 10 most abundant AC subtypes identified in the recent scRNA-seq have no previously established marker and have not been studied in any way (Yan et al., 2020). In addition, ACs also have an extra layer of laminar complexity due to their presence in both the INL and the GCL. Previous studies of a limited number of AC subtypes suggest that the broader AC classification of GABAergic or glycinergic receptor expression have distinct sublayer distribution (Cherry et al., 2009; Voinescu et al., 2009). Our findings are generally consistent with previous reports, where most GABAergic subtypes were found in the basal INL and glycinergic subtypes were found in the central INL. Of note, 3 nGnG subtypes, which share transcriptional similarity with glycinergic subtypes, were among the most apically positioned AC subtypes in the INL. The three nGnG uniquely express a transcription factor Ebf, and our observation is supported by the previous immunohistostaining result of Ebf+ GlyT1- cells in the most apical sublayer of ACs in INL (Kay et al., 2011; Voinescu et al., 2009).

The mechanism underlying the highly regulated process of establishing laminar architecture in the retina has been previously investigated (Amini et al., 2018). On the major cell type level, there is a correlation between the birth date and the layer position, of which cells born early take a more basal position (Clark et al., 2019). There are exceptions to this rule such as MGs, which are the last cell type to form and are located in between ACs and BCs. On the subtype level, similar relationship between the birth date and the layer position can also be observed. For example, previous studies that have demonstrated distinct sublayer distribution of GABAergic and glycinergic AC subtypes have also described a linear relationship between the birth order and basal-to-apical nuclei positioning. (Cherry et al., 2009; Voinescu et al., 2009).

We similarly observed a general basal-to-apical pattern in the subtype genesis with earlier born subtypes (GABAergic) distributed between the GCL and the basal INL and the later born subtypes (glycinergic) in the apical INL. In contrast, the relative position of ON and OFF BCs was less correlated with their birthdate. Although it is known that OFF BC subtypes are born before ON BC subtypes (West et al., 2022), we observed significant mixing between OFF and ON BC positioning and did not find any difference in the averaged radial position between the two groups. Moreover, while RBCs are born between OFF and ON BCs, we found that RBCs were located at the most apical layer in the INL. Thus, our observation suggests that a significant adjustment of cell positioning occurs at both major cell type, subgroup, and subtype level during development.

#### Clone mosaic pattern

The examination of how neuronal subtypes are dispersed in the corresponding laminar sublayers revealed that the enrichment or depletion of direct neighboring subtype pairs of any given neuronal subtype is rare, and the distribution of subtypes is mostly random. The general randomness in direct neighbor relationship is consistent with the idea that retinal neurons spread out to maximize tissue coverage and the cells of same subtypes tend to be away from each other. A few enriched or depleted homotypic and heterotypic subtype pairs in our data suggest that there may be complex homophylic or repulsion mechanism in the arrangement of neuronal subtypes in the retinal maturation. As a recent study of the BC subtype genesis demonstrated that the postnatal clones are biased against homotypic BC subtypes (West et al., 2022), our finding of enriched BC9 homotypic pairs could be explained by the large number of cells profiled in our experiment. While the retinal cell types are widely believed to be assembled in regular soma arrays in retinal mosaics, our data revealed that almost all neuronal subtype soma are dispersed randomly with no more significant regularities compared with random simulations, contradicting a common assumption. This finding indicates that the cell distribution of retina is achieved at the level of dendritic field patterning rather than the neuronal soma placement.

### Cell location provides an important dimension for its function

A total of twelve AC subtypes with significant displacement patterns were identified in our study. Among them, eight subtypes (AC2, AC7, AC21, AC32, AC39, AC44, AC46, and AC57) are novel subtypes without any established key marker prior to the scRNA-seq study (Yan et al., 2020). Interestingly, all displaced subtypes were GABAergic except for AC36, which is a nGnG type. This is consistent with the idea that most displaced ACs are born early and occupy a more distal position. The AC subtypes that are not preferentially displaced accounted for 29% of total ACs in the GCL. It is not clear whether the displacement of ACs occurs due to the functional necessity exemplified by their axon strata or simple mis-migration during development. One possibility is that the 29% of amacrine cells we identified in the GCL with no statistical displacement may truly have their cell bodies mis-localized in the GCL.

The starburst AC (SAC) is one of the most studied displaced AC subtypes. Although the two groups of SACs located in the INL and the GCL cannot be distinguished based on current scRNA-seq, they are considered two subtypes and possess distinct functional differences derived during final cell migration and dendrite formation (Tauchi and Masland, 1985; Vaney, 1990). To our surprise, DEG analysis of the SACs in two different locations identified a set of differentially expressed genes, including several known ones (Peng et al., 2020). As exemplified in the case of displaced ACs, the cell identity and function are determined by the combination of the unique transcriptome as well as cell location. Our finding illustrates the limitation of identifying cell types based on scRAN-seq alone and the importance of including other information, such as the spatial location of the cells, in achieving a comprehensive list of cell types in the tissue.

## Conclusion

We have generated the first spatial atlas of the mouse retina using spatial transcriptomics and integration with scRNA-seq. Our data allowed systematic evaluation of near all known cell subtypes and revealed interesting aspects of cell organization in the retina. Laminar organization of cells was observed at the major cell type and subgroup level, both of which have strong correlation with the birth order of cells. Random dispersion of neuronal subtypes was observed, contradicting the common assumption of retinal neurons are assembled as regular soma array. In conclusion, the spatial atlas can serve as a powerful resource for the community and provide the foundation for further investigation on the interplay among cell identities, locations, and functions.

## Acknowledgements

This project was funded by NIH/NEI R01EY022356, R01EY018571, S10OD032189, Chan Zuckerberg Initiative (CZI) award CZF2019-002425, RRF to R.C, and NLM fellowship program T15LM007093 to S.F. J.R.M is a founder of, stakeholder in, and scientific advisor for Vizgen. J.R.M is an inventor on patents associated with MERFISH applied for on his behalf by Harvard University and Boston Children’s Hospital and licensed to Vizgen. The other authors declare no competing financial interests. We thank Vizgen for the transcript probes, the transcript decoding pipeline and the experimental guidance.

## Author contributions

Experiment conceptualization J.C, J.L, S.F, Q.L and R.C; MERFISH experiment J.C, and S.F; Data analysis: J.C and J.L; Data resource development J.L; RNAScope validation J.C; Writing, reviewing and editing J.C, J.L, S.F, J.R.M, and R.C.

## Methods

Animal Studies: Mouse housing, experiments, and handling were approved by the Baylor College of Medicine Institutional Animal Care and Use Committee, and the studies were conducted in adherence with the ARVO Statement for the Use of Animals in Ophthalmic and Vision Research and followed the guidance and principles of the Association for Assessment and Accreditation of Laboratory Animal Care. C57Bl/6J mice were bred in-house and maintained in a 14h light / 10h dark cyclic environment. Animals were housed by Baylor College of Medicine Center of Comparative Medicine.

Sample Collection: Whole eyes were enucleated and immediately flash frozen on dry ice after being embedded in TissueTEK O.C.T. compound (VWR Cat No. 25608-930). Samples were then stored in -80°C prior to MERFISH or RNAScope experiments. Experiments were performed using about P90 mice.

MERFISH probe panel design: In designing the probe panel list, we included a handful of major cell type markers for rod photoreceptors, cone photoreceptors, BCs, ACs, RGCs, HCs, and MGs. In addition, several transcription factors critical for the retina tissue function were included. To capture the cell heterogeneity on the subtype level, we included further 3 to 4 top-ranked genes of each BC subtype cluster and 1-2 top-ranked genes of each AC and RGC subtype cluster identified in the publicly available scRNA-seq data (Shekhar et al., 2016; Tran et al., 2019; Yan et al., 2020). The heatmap of the probe transcript expression was plotted using a down-sampled reference data containing 1,000 of rod photoreceptors, cone photoreceptors, BCs, ACs, RGCs, MGs, and 359 HCs.

Multiplexed Error-Robust Fluorescence in situ Hybridization (MERFISH): This protocol was adapted for fresh frozen retinas by Vizgen. In brief, 20mm functionalized coverslips (Vizgen, #FCS01) were treated with 1% polyethyleneimine for 1 hour at room temperature and washed with nuclease-free water. Coverslips were then coated with yellow green fluorescent fiducial beads (Polysciences, 17149-10) in PBS for 10 minutes at room temperature. The coverslip was washed twice briefly with nuclease free water and allowed to air dry. Afterwards, 10µm cryo- sections were cut near the vicinity of the optic nerve and placed onto the bead coated surface of the coverslip. The tissue sections were fixed at -20°C with pre-chilled 100% ethanol for 30 minutes. The sample was brought to the benchtop and permeabilized in fresh room temperature 100% ethanol for 1 hour. The sample was re-hydrated by sequentially exchanging buffers to 90% and 70% ethanol in a 5-minute interval and washed briefly in 2X SSC buffer. The sample was incubated for 30 minutes at 37°C in 5mL of formamide buffer before 50µL of a custom designed probe panel containing 22 bits (Vizgen) was placed directly onto the retinal tissue. The sample was incubated with the probes in a 37°C cell culture incubator for 36-48 hours (about 2 days).

After probe hybridization was complete, the sample was washed twice with formamide buffer for 30 minutes each at 47°C and then briefly with 2X SSC three times to remove any residual formamide buffer. Cell membrane staining was done using oligo-conjugated primary and secondary antibodies provided by Vizgen (Vizgen, #CB-MM). In brief, the tissue is blocked for 1-hour at room temperature with 105µL total volume of blocking solution (100µL of blocking buffer plus 5µL of murine RNAse inhibitor). Then a 1:100 primary antibody dilution was made in the blocking solution and incubated on the tissue for 1 hour at room temperature. The sample was washed three times with PBST for 5 minutes each on a rocker and then a 1:33 secondary antibody solution was incubated on the tissue for 1 hour at room temperature. The sample was then washed three times with PBST for 5 minutes each on a rocker and then three times with 2X SSC. Afterwards, the sample was incubated in 50µL of an acrylamide/bis-acrylamide based gel solution for 1.5 hours at room temperature before being cleared for overnight or until the tissue becomes transparent at 37°C. After tissue clearing, the sample was washed several times in 2X SSC before the initial hybridization of fluorescence readout probes against the oligo-conjugated antibodies for for 15 minutes at room temperature. Lastly, the sample was incubated in the wash buffer from the Vizgen Imaging Reagent Kit (Vizgen, #IK-24) for 10 minutes at room temperature before sequential barcode images were imaged on the Vizgen Alpha Instrument.

Sequential extinguishing of the fluorescent signal and re-hybridization fluorescent read-out probes were performed by the automatic fluidic system on the Vizgen Alpha Instrument.

### Data analysis

MERlin decoding: Raw MERFISH images were decoded by the MERlin pipeline (v0.1.9, provided by Vizgen) (Emanuel et al., 2020) using the 22-bit codebook designed for 368 genes in the MERFISH panel. Following MERlin tasks were perfomed: FiducialCorrelationWarp by merlin.analysis.warp module; DeconvolutionPreprocessGuo by merlin.analysis.preprocess module with warp_task: FiducialCorrelationWarp, highpass_sigma:2, decon_sigma:1.4, and decon_filter_size:9; ten rounds of Optimization by merlin.analysis.optimize module with area_threshold: 5 and fov_per_iteration: 50; CorrelationGlobalAlignment by merlin.analysis.corralign module with crop_width:100; Decode by merlin.analysis.decode module with preprocess_task: DeconvolutionPreprocessGuo, optimize_task: Optimize10, global_align_task: CorrelationGlobalAlignment, minimum_area: 1, lowpass_sigma: 0.6, crop_width: 100, write_decoded_images: false, distance_threshold: 0.6, remove_z_duplicated_barcodes: true; GenerateAdaptiveThreshold by merlin.analysis.filterbarcodes module with decode_task: Decode, run_after_task: Optimize10; AdaptiveFilterBarcodes by merlin.analysis.filterbarcodes module with decode_task: Decode, adaptive_task: GenerateAdaptiveThreshold and misidentification_rate: 0.05; ExportBarcodes by merlin.analysis.exportbarcodes module with filter_task: AdaptiveFilterBarcodes, columns: [“barcode_id”, “global_x”, “global_y”, “global_z”, “x”, “y”, “fov”]. Decoded transcripts, which represent single pixels, were exported with their barcode ID and coordinates in csv file format.

Cellpose and Mesmer cell segmentation: To detect cell boundaries in MERFISH images, cell segmentation was performed on cell boundary staining and DAPI images of individual field-of- views (FOVs). To increase relative brightness and darkness of cell membrane and DAPI staining from the background, images were adjusted by contrast enhancement using CLAHE Histogram Equalization function in OpenCV (Bradski, 2000). To increase the number of segmented cells per FOV, consecutive stacks Z0 and Z1 were blended by averaging intensities of the two stacks. Using the blended cell membrane images, Cellpose (v0.6.5) (Stringer et al., 2021) was used to predict cell boundaries using the “cyto2” mode, which was trained on a larger dataset submitted by users. Due to the difference in compactness of the three nuclear layers that results in distinct background and staining intensities, the Cellpose generated definitive segmentation result in the ONL and INL, but not in the GCL. This may be attributed to the high signal intensity coming from the dense extracellular matrix in the GCL. To rescue non-segmented cells particularly in the GCL, we applied Mesmer (Greenwald et al., 2022) using “nuclear” for compartment parameter and 0.1667 for image_mpp parameter using 2-stack cell membrane and DAPI images.

Mesmer resulted in near complete segmentation in the GCL, but often showed high doublet detection rate in the ONL and INL. To achieve optimal segmentation in all three retinal layers, segmented cells from Mesmer were retained if they overlapped <0.1% area of cell polygons by Cellpose. Decoded transcripts were then assigned to segmented cells by searching nearest polygons using the k-d tree algorithm in the SciPy (Virtanen et al., 2020). To retain high-quality cells, segmented cells were further filtered by average DAPI intensities (>= 80), radius of minimum enclosing circles (>= 10, <= 80), area of polygons (>=500, <=10,000), perimeter of polygons (>=50, <=400), and total transcripts (>= 10).

Integration between MERFISH experiments: To reduce the batch effect between MERFISH experiments, batch correction was done to integrate different samples using scVI (Lopez et al., 2018), which models the gene expression on a batch variable as well as library size and latent representation. To perform the scVI integration, raw counts of full 368 genes in the MERFISH panel were used with the four “sampleID”s as the batch variable with 2 hidden layers for encoder and decoder neural networks and 30 dimensionality of the latent space. The generated 30 dimensional scVI low-representations were used to calculate a 2D UMAP for visualization of MERFISH cells by Scanpy (Wolf et al., 2018). The low-representations were also used to measure dissimilarities among cells, and the dissimilarities were used to calculate the cell clustering by the Leiden algorithm (Traag et al., 2019) with resolution 1.

scRNA-seq meta analysis: scRNA-seq meta analysis: The scRNA-seq reference used in subtype annotation was generated by combining the publicly available scRNA-seq data for BC, AC, and RGC (Shekhar et al., 2016; Tran et al., 2019; Yan et al., 2020) and our in-house single-cell/nuclei RNA-seq data. Our in-house data, which have not been published, are composed of about 130,000 cells, in which 14,000 ACs, 70,000 BCs, 400 RGCs are included. Collected datasets underwent a standardized preprocessing to exclude empty droplets, ambient RNAs, and estimated doublets (https://github.com/lijinbio/cellqc). Retained data were integrated using scVI (Lopez et al., 2018) to reduce the batch effect. The trained low-dimensional representations were used to calculate the dissimilarities among cells, and cell clusters were detected by the Leiden algorithm (Traag et al., 2019). To annotate the major cell types, top ranked genes were calculated compared to the rest of cell clusters and used to match major cell type markers. Similarly, for AC, BC, and RGC-subtyping, data integration and cell clustering were performed on corresponding subtypes, respectively. Published subtype labeling was retained and used to guide the annotation of subtype cell clusters. BC and RGC-subtyping showed exhausted subtypes.

After integrating our in-house data, over-clustering of some AC subtypes annotated in public datasets were merged into single cell clusters. To retain the public subtype labels, the merged cell clusters were labelled by the subtype of the majority number of cells. For example, a merged cell cluster was labeled as AC21 while it includes subtypes AC21, AC54, and AC63 in the public datasets.

Cell identity assignment: To identify the cell type of MERFISH cells, a two-level annotation was performed for major cell type annotation and subtype annotation in sequence. To annotate the major cell type of MERFISH cells, we first calculated top ranked genes of Leiden cell clusters using Scanpy and manually inspected the known major cell type marker genes to assign major cell type identities (Traag et al., 2019; Wolf et al., 2018). Following the major cell type annotation, subtype annotation was performed by co-embedding method using scRNA-seq reference data for isolated BC, AC, and RGC subset. Using bindSC (Dou et al., 2022), scRNA-seq reference and MERFISH count matrices were bridged by optimizing correlation in the sample and feature levels simultaneously. Using bindSC, which integrates two different single-cell modalities by optimizing correlations in the sample and feature levels simultaneously, the reference scRNA-seq and MERFISH count matrices were aligned. The generated canonical correlation vectors are co-embedding low-dimensional (e.g., 15) latent representations for scRNA-Seq and MERFISH cells. These co-embeddings were used to calculate a 2D UMAP between the two modalities. To annotate subtypes of MERFISH cells, the cell labeling of scRNA-seq reference was transferred by SVM classifiers. Specifically, the latent representations of scRNA-seq cells were used to train multi-class SVM classifiers by the known scRNA-seq cell labels. Per MERFISH cell, scRNA-Seq neighbors (e.g., 3 neighbors) were detected in the co-embedding latent space, and the average of low-dimensional representations was used to represent the feature for the classification. Classification probabilities were calculated against the trained SVM classifiers, and the cell type was assigned by the maximum classification probability of the classifiers. BC, AC and RGC subtypes are annotated by applying the label transfer using the bindSC co-embedding.

Cell clustering using MERFISH genes and subtype annotation overlay: Subtype annotation from co-embedding analysis using reference scRNA-seq data was overlaid on the graphical clustering map derived from using only MERFISH transcriptome. Subsets of BCs, ACs, and RGCs were extracted from a representative MERFISH experiment (“sampleID” = “merfish_wt_JC2”).

Leiden clustering was performed using Scanpy in each cell type data subset (Traag et al., 2019; Wolf et al., 2018), and cells were colored by the subtype annotation labels from bingSC co-embedding.

Tangram imputation: To resolve the entire transcriptome of MERFISH cells, we impute the gene expression by mapping single-cell RNA-Seq reference to spatial cells using Tangram (Biancalani et al., 2021). Tangram calculates the imputation by summing over scRNA-seq gene expression weighted by mapping probabilities. Tangram trains the mapping probabilities by optimizing correlations between imputed expressions and MERFISH measurements of the MERFISH panel. To reduce the bias caused by scRNA-seq cells with a different cell type from a MERFISH cell, mapping probabilities of the only scRNA-seq cells with the same cell type were used for the imputation. To minimize the crossover and retain the spatial proximities of subtypes in the co-embedded space, imputation was performed within isolated AC, BC, and RGC subtypes separately with the subtype labeling annotated by bindSC co-embedding. To measure the performance of Tangram imputation, Pearson correlation coefficients have been calculated between imputed gene expressions and MERFISH measurements for the genes included in the MERFISH probe panel. A UMAP (McInnes et al., 2018) was also generated using imputed gene expressions to visualize the labeling of MERFISH cells.

Boundary estimation and distance calculation: The ONL and INL boundaries were estimated by approximating surrounding curves that cover the centroids of segmented cells in micrometers. To do this, the R package *alphahull* (Pateiro-López and Casal, 2010) was used to calculate the alpha-shape and alpha-convex hull of cell centroids. Alpha-shape was defined as an extension of convex hull, where the shape was segments connecting alpha-extreme points. A cell was called an alpha-extreme point when an open ball of radius alpha existed with the centroid on the boundary and did not cover other cells. The alpha-convex hull stored the arcs of open balls that connect neighboring alpha-extreme points. The alpha value of 100 micrometers generated the alpha-shape with a tight estimate of tissue boundaries including basal and apical boundaries. To separate basal and apical boundaries, the nature of an arc-shape for a retina tissue was utilized. Each section of retina tissue was assumed to have a hypothetical center. This tissue center was compared with each arc of the alpha-convex hull. Apical boundaries contained arcs that were outward away from the tissue center. On the contrary, arcs of basal boundaries were inward to the tissue center. A small number of mis-labeled boundaries were corrected interactively using the R package *shiny* (Chang et al., 2021). To calculate distances of cells to the two estimated boundaries, perpendicular distances were calculated to each segment of boundaries using the function *nearestPointOnSegment()* in the R package *maptools* (Bivand et al., 2022). The minimum distance of segments was the calculated distance of a cell to a boundary. To normalize the thickness among tissue sections, the distance ratio was calculated by the distance to the basal distance over the sum of distances to the two boundaries.

Determining significance of cell displacement: To identify AC subtypes displaced in the GCL, the INL boundary information previously generated was used to assign ACs to their resident layer. For each subtype, the proportion of cells displaced in the GCL was calculated. To determine the significance of displacement, we applied a permutation test to calculate the expected proportion of displaced ACs by shuffling the subtype label within tissue sections for 1,000 times (Casella and Berger, 2021). The *p-*value was calculated as the percentage of permutations that have a greater proportion of cells in the GCL layers compared to the observed proportion. The AC subtypes with *p*-value < 0.05 were determined to be significantly displaced.

RNAScope in-situ hybridization: The RNAscope HiPlex Assay (ACD Biosystems) (Wang et al., 2012) was performed according to the ACD protocol for fresh-frozen tissue. In brief, whole mouse eyes embedded in TissueTEK O.C.T. compounds were cut in 10µm sections and placed on glass microscope slides. After fixation with 4% FPA in PBS and permeabilization using Protease IV (RNAScope), the tissue sections were incubated with probes against displaced AC markers, pan-AC marker Slc32a1 (319191-T1), and pan-RGC marker Slc17a6 (319171-T2).

Following probes were to stain for specific displaced AC subtypes; AC17 – Slc18a3 (448771- T5), AC21 (495681-T7), AC32 (402021-T9), AC36 (834921-T10), AC37 (316091-T3), AC39 (421011-T6), AC44 (317451-T11), AC46 (452981-T5), AC48 (413561-T6), AC52 (1142431-T7). The probes were amplified according to the manufacturer’s instructions and labeled with the following fluorophores for each experiment: Alexa 488 nm, Atto 550 nm, and Atto 647 nm. The Zeiss Apotome.2 microscope (Zeiss Axio Imager) was used to visualize the FISH signals.

Nearest neighbor relationship analysis: To investigate the nearest neighbor relationship, the R package *Giotto*(Dries et al., 2021b) was used to construct a pairwise cell-cell spatial network through the Delaunay triangulation method. The function *createSpatialNetwork()* was used to build the Delaunay network with the maximum distance of 14 microns for cells in the INL and 20 microns for cells in the GCL. The maximum distance used was based on the cell-to-cell distance measured from immunohistochemistry images. To integrate tissue sections, cells from individual sections were stitched into one image by shifting each section by 4,000 µm in both axes. After visual inspection of the created network, cell proximity scores, which represents an enrichment or depletion of pairs between cell types, were determined by calculating the observed over expected frequency of cell-cell interaction pairs. The expected frequency was simulated through label-shuffling of cell type in fixed cell coordinates. To compensate for the laminar structure of retinal cells, the label-shuffling was performed within 10 bins of the normalized INL length. Using the *cellProximityEnrichment()* function, cell proximity scores within the three subsets of population, which contain BCs only, ACs in the INL only, and both ACs and RGCs in the GCL, were calculated via 1,000 simulation, and the p values were adjusted using false discovery rate. The enrichment scores and p values included in the data table outputted by *cellProximityEnrichment* was used to generate the heatmap using *ggplot* library in R.

Nearest neighbor regularity index analysis: To examine the interspacing pattern of neuronal subtypes in the retina, a nearest neighbor regularity index (NNRI) was calculated per each subtype(Keeley et al., 2020). The NNRI was calculated by determining the average nearest neighbor distances for individual cells belonging to the same subtype across clean tissue sections of wild-type samples over the interquartile range (IQR) of the distance distribution. The statistical significance is calculated by a permutation test by label shuffling within each tissue section for 1000 times, and NNRIs are calculated per subtype for each permutation. To compensate for the laminar structure of retinal cells, the label shuffling was performed within 10 bins of the normalized INL length. Each of 1000 NNRIs from permutations was used as a background distribution for subtypes in the histogram compared with observed NNRI in the red vertical line. The *p*-value was determined as the percentage of NNRIs from permutation that is greater than that of the observed NNRI. *p*-value < 0.05 or *p*-value > 0.95 was used to identify the non-random subtypes. To characterize RGCs and ACs in the monolayer of GCL layer, the label shuffling was performed in the one layer of GCL. *ggplot* library in R was used to generate the histogram.

Differential gene expression analysis by location: To identify the differentially expressed genes between ACs in the GCL and ACs in the INL, we adapted the workflow of *DESeq2 (Love et al., 2014)* for the imputed gene expression. First, the imputed gene counts were normalized using the function *computeSumFactors()* in the R package *scran*. Then, the normalized values were fit into a negative binomial model by *glmGamPoi*. The likelihood-ratio test was applied to test the statistical significance by comparing with a reduced model *∼1*, and the calculated *p*-values were adjusted by the Benjamini–Hochberg procedure. The log2 fold changes were also reported in the result table. Differentially expressed genes were identified under *q*-value<0.05 and |log2FoldChange|>0.2. To visualize the DEGs, the volcano plot was generated by the R package *EnhancedVolcano* (Blighe et al., 2018).

Data resource visualization: Access to the spatial map of mouse retina is hosted at https://bcm.box.com/s/bwh011jg0m7b6t38j3q6chsvf9xbk0wg. The data is shared in a .h5ad format with gene expression matrix and curated metadata. To provide a user-friendly interactive exploration of the spatial map, the imputed gene expression are made accessible via cellxgene (Megill et al., 2021). The two plots contained in the data resource; the tissue plot showing the coordinates of identified MERFISH cell centers and UMAP visualization graph calculated using the imputed gene expressions. The metadata of annotated cell types can be used to color the cells in the plots. Furthermore, expression pattern of a gene or set of genes can be visualized in a histogram and in the tissue plot with the color scale as expression value. The URL of the cellxgene web service is accessible at http://cellatlas.research.bcm.edu.

Data and Software availability: The data in this study are available from the corresponding author on reasonable requests. MERFISH data are shared via the data portal at Baylor College of Medicine. The MERFISH count matrix in the .h5ad format with metadata such as annotated cell type label and coordinates can be accessed at https://bcm.box.com/s/bwh011jg0m7b6t38j3q6chsvf9xbk0wg. The code for MERFISH image analysis is available at https://github.com/RCHENLAB/SpatialMmMERFISH.

## Supplemental Figures

**Supplemental 1.**
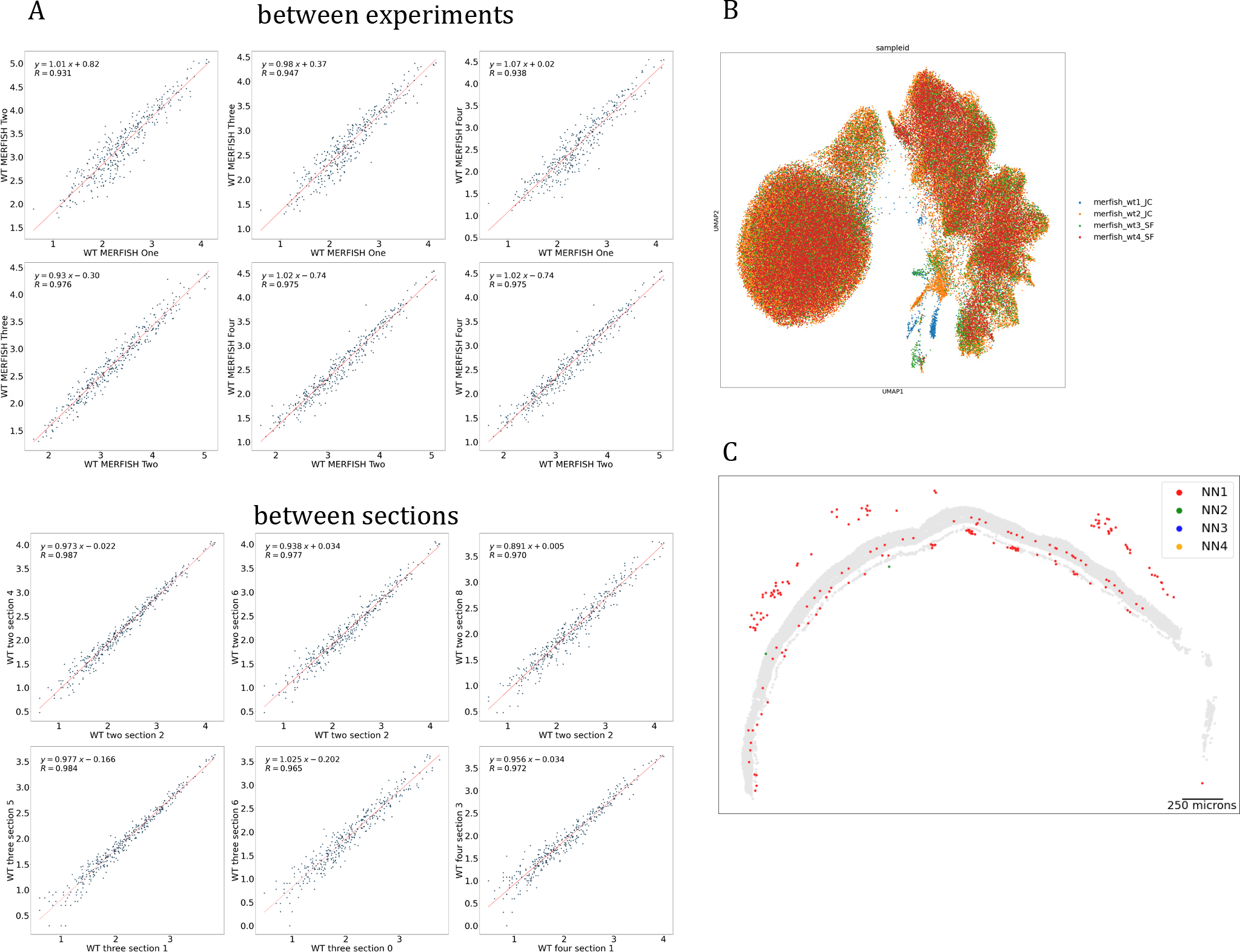
MERFISH data quality and integrative clustering results of 4 MERFISH experiments. **(A)** Pearson correlation coefficients between four MERFISH experiments and between retina tissue sections within the same experiment. High reproducibility can be observed between tissue sections and experimental replicates. **(B)** Integrative clustering analysis of MERFISH experiments. Batch correction between four MERFISH experiments resulted in even distribution of cells with strong overlap. We observed cell clusters that are batch specific with no unique expression of transcripts. These clusters were subsequently determined to be non-neuronal cells. **(C)** Distribution of non-neuronal retinal cell types. Non-neuronal cell types cannot be accurately assigned to a cell type due to the lack of specific markers in the panel. The cell coordinates and gene expression of these cells suggest these are astrocytes, RPE, and immune cells.

**Supplemental 2.**
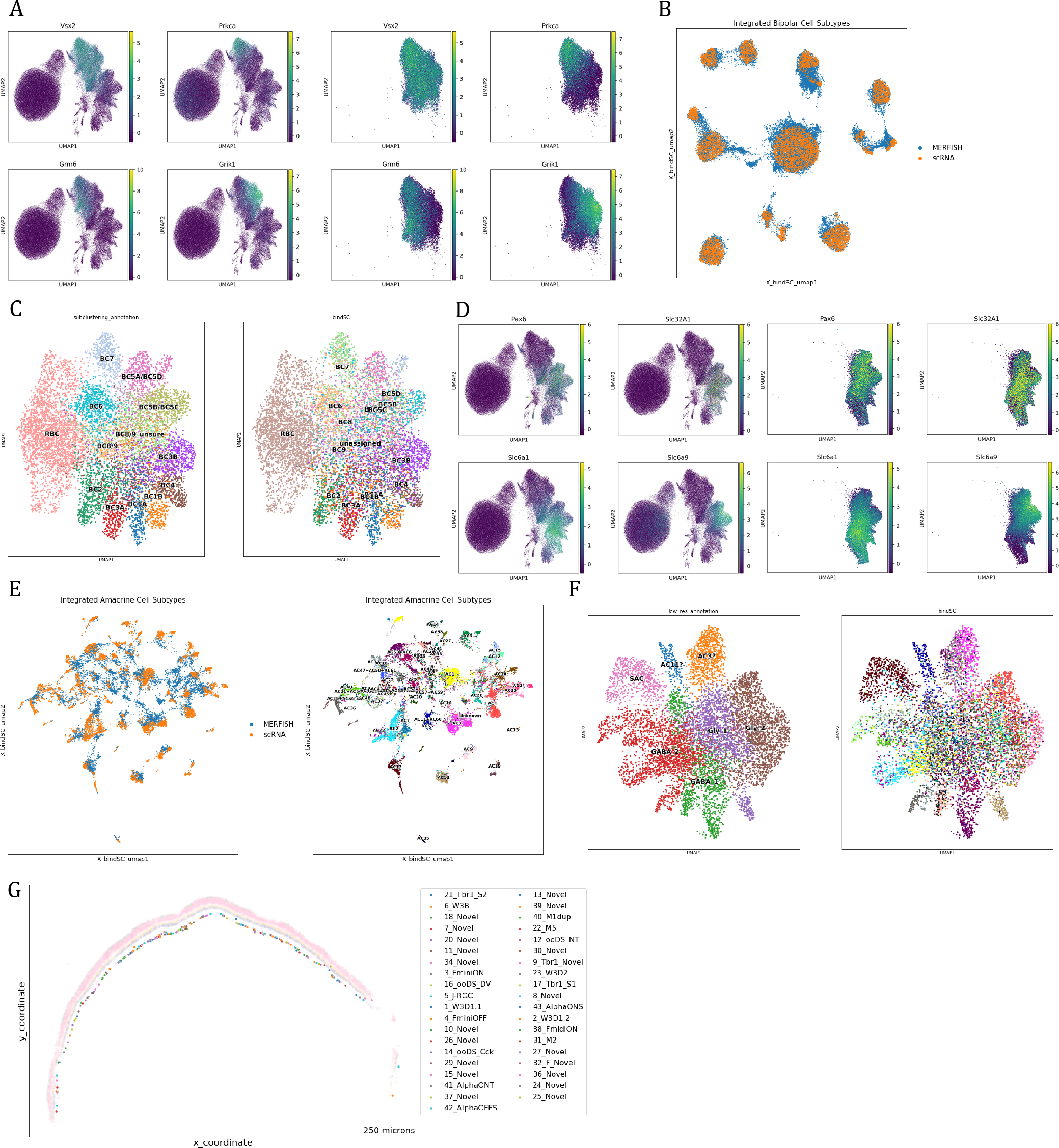
Broader classification of BC and AC types, and co-embedding analysis quality check. **(A)** Bipolar subgroup marker expression. A pan-BC marker Vxs2 is expressed across all BCs. BC subgroup markers such as Grm6 for ON cone BC, Grik1 for OFF cone BC, and Prkca for RBC are expressed specifically within the substructures of BC clusters. **(B)** BC subtype label overlaid on the lower dimensional space calculated using only 368 MERFISH features. Clustering analysis with only the MERFISH features results in poor resolution, as shown in clustering map with merged subtypes on the left. Overlaying the subtype annotation from co-embedding analysis on the same map shows separation within previously merged subtypes such as BC5A/BC5D in the right plot. **(C)** Co-embedding plot of BC from MERFISH and scRNA-seq. BCs identified in MERFISH (blue) and scRNA-seq (orange) show strong overlap. **(D)** Amacrine subgroup marker expression. Pan-AC markers Pax6 and Slc32a1 are broadly expressed across all ACs. Expression of GABA and glycinergic receptor markers, Slc6a1 and Slc6a9 was confined within substructures of AC clusters. **(E)** Co-embedding plot of amacrine cells from MERFISH and scRNA-seq. ACs identified in MERFISH (blue) and scRNA- seq (orange) show reasonable overlap. **(F)** AC subtype label overlaid on the UMAP with 368 MERFISH features. Clustering analysis using the MERFISH features does not provide sufficient resolution to identify AC subtypes as shown in the left plot. The co-embedding annotation on the same UMAP on the right shows that AC subtypes are confined within substructures of clusters, represented by clumped up cells in same colors. **(G)** Tissue plot of identified retinal ganglion cell subtypes. All RGCs are seen in the GCL, where they are known to reside.

**Supplemental 3.**
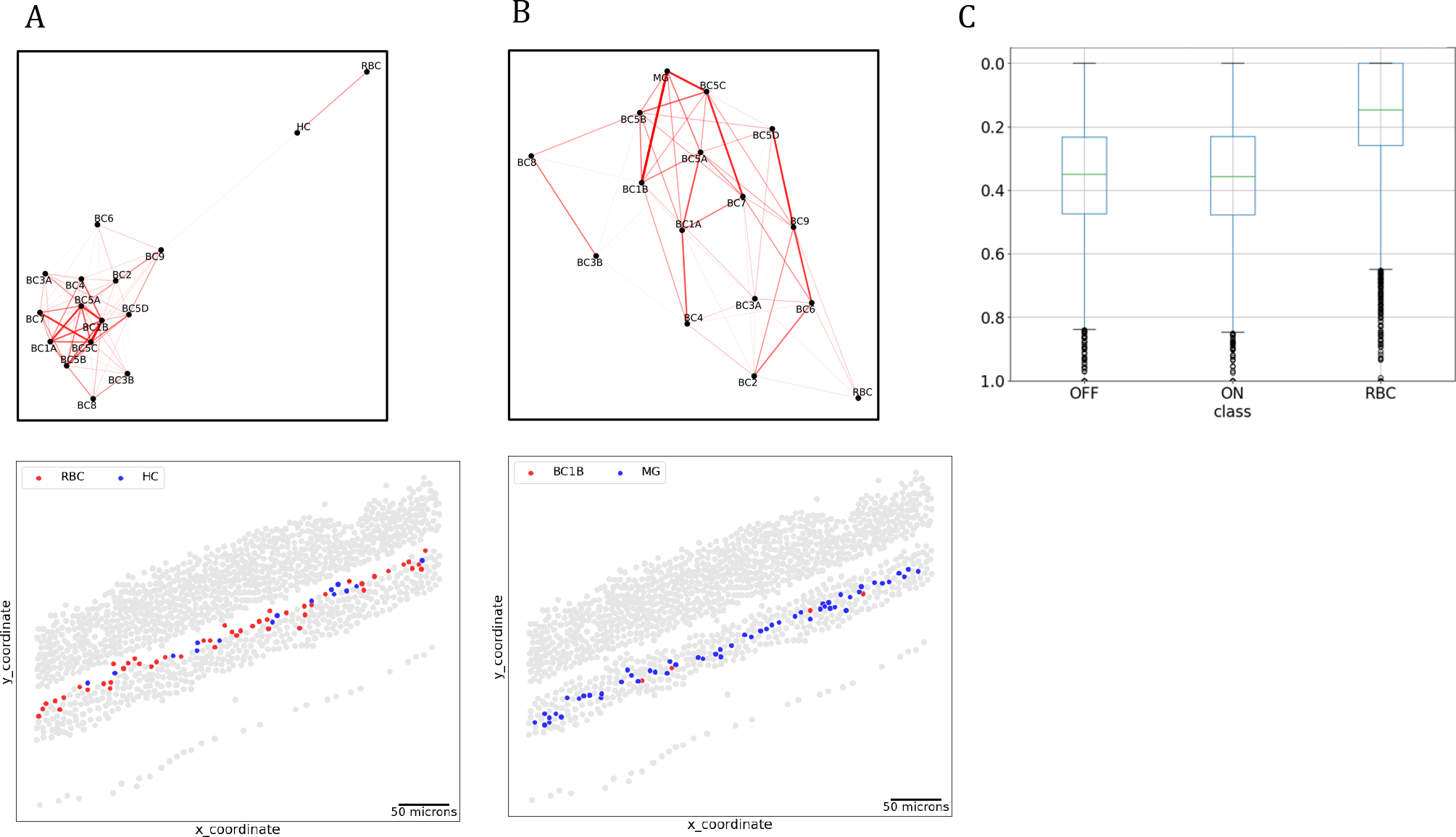
Spatial relationships of RBC and BC1B, and average INL position of OFF, ON, and rod bipolar types. **(A)** Spatial relationship between rod bipolar cells and horizontal cells. RBC and HC show high proximity enrichment score as shown in the network plot on top. The tissue plot in the bottom indicates both RBC and HC are positioned in the apical layer of the INL. **(B)** Spatial relationship between BC1B and Muller Glial cells. Network plot on top show high proximity enrichment score for BC1B and MG The tissue plot in the bottom indicates BC1B are positioned in the central INL along MG. **(C)** Average distance of OFF cone, ON cone, and RBC in the normalized INL. No significant difference between OFF and ON cone BC distance was found. RBCs show significant apical positioning compared with cone BCs.

**Supplemental 4.**
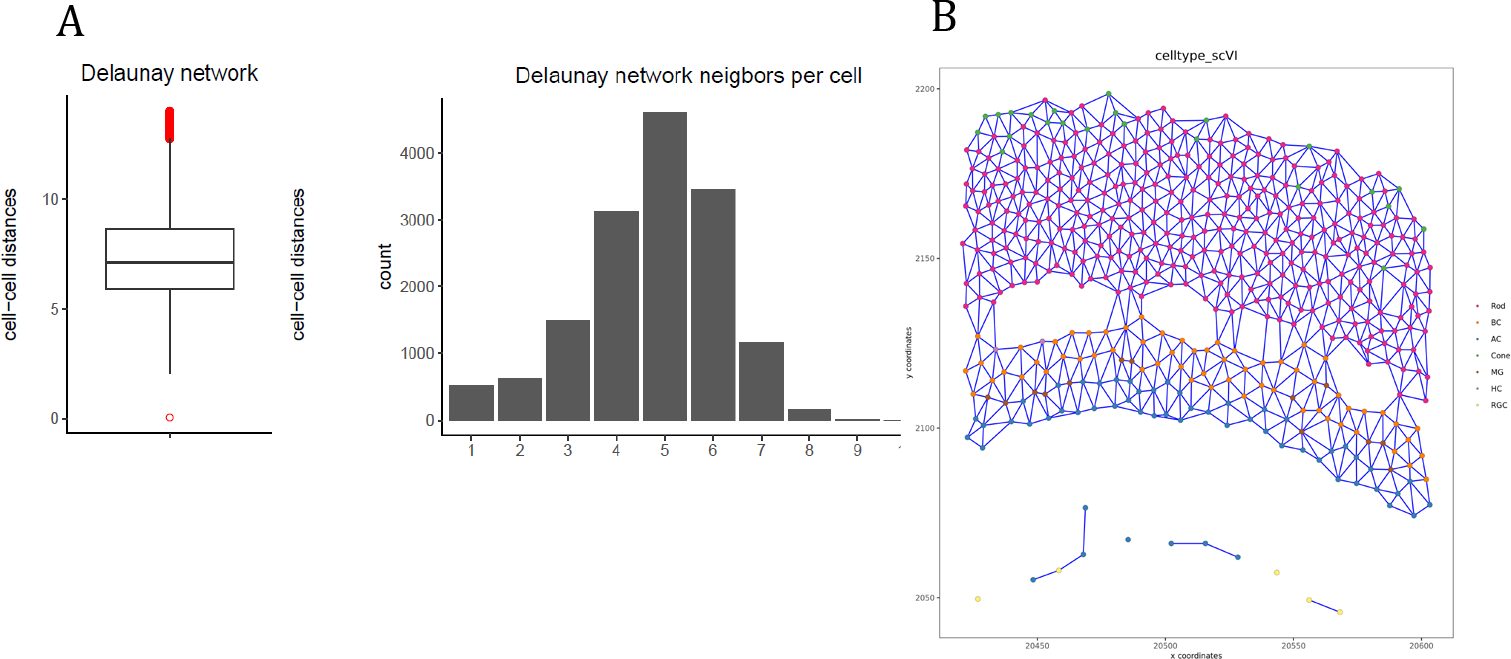
Pairwise cell-cell network statistics. **(A)** Delaunay triangulation method with maximum distance of 14 µm between each cell was used to construct the network. Cells have about 5 nearest neighbors with average distance of ∼7 µm, indicating that the network mostly consisted of immediately adjacent cells. **(B)** An example plot of cells and their network connection is shown.

**Supplemental 5.**
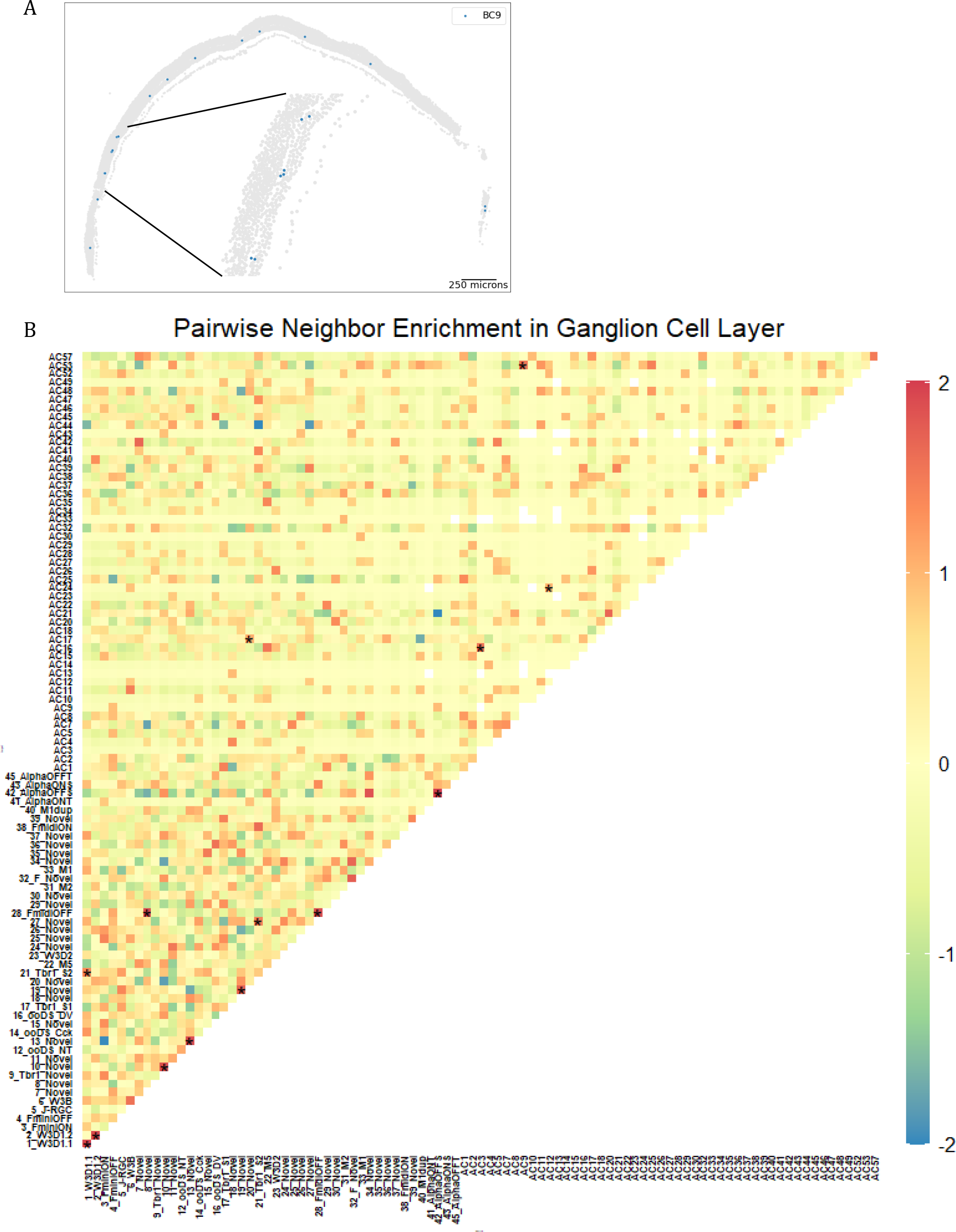
BC9 homotypic pairs, and nearest neighbor relationship in GCL. **(A).**Example plot showing homotypic pairs of BC9 **(B).** Heatmap of pairwise nearest neighbor enrichment scores of RGCs and ACs in the GCL. Pairwise nearest neighbor scores within cells in the GCL were calculated to determine any significant subtype pairs. Displaced ACs and RGCs show random distribution patterns with a few exceptions. Enriched or depleted nearest neighbor pairs with statistical significance are labeled with asterisks.

**Supplemental 6.**
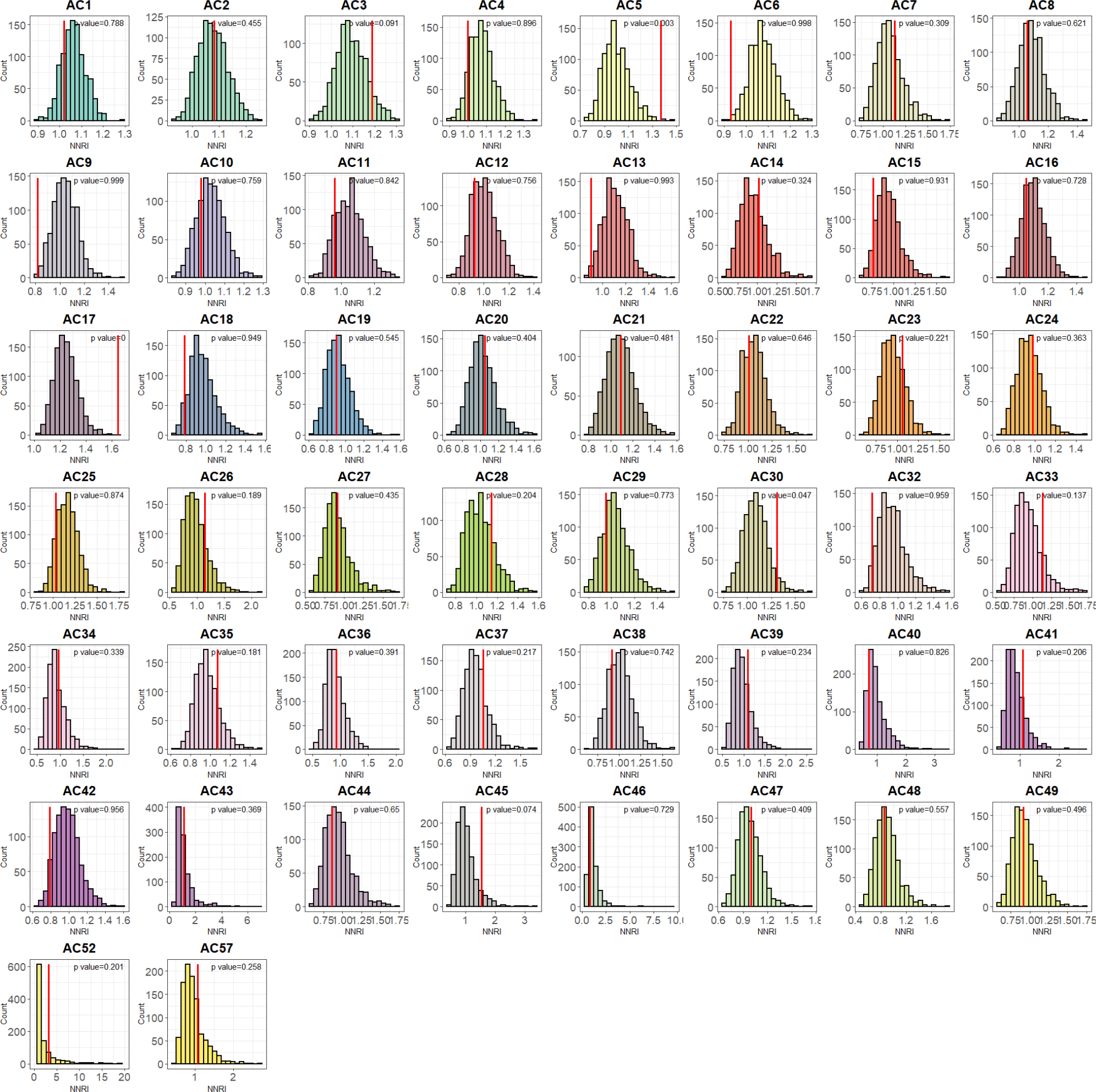
Comparison between observed and random regularity indexes of amacrine cell subtypes. All except for six AC subtypes showed non-significant difference in NNRI (in red line) compared with random permutation shown in the histogram. AC6, AC9, and AC13 showed smaller NNRI compared with random permutation, indicating shorter distance between cells. AC5, AC17, and AC30 showed larger NNRI (in red line) compared with random permutation, indicating larger distance between cells.

**Supplemental 7.**
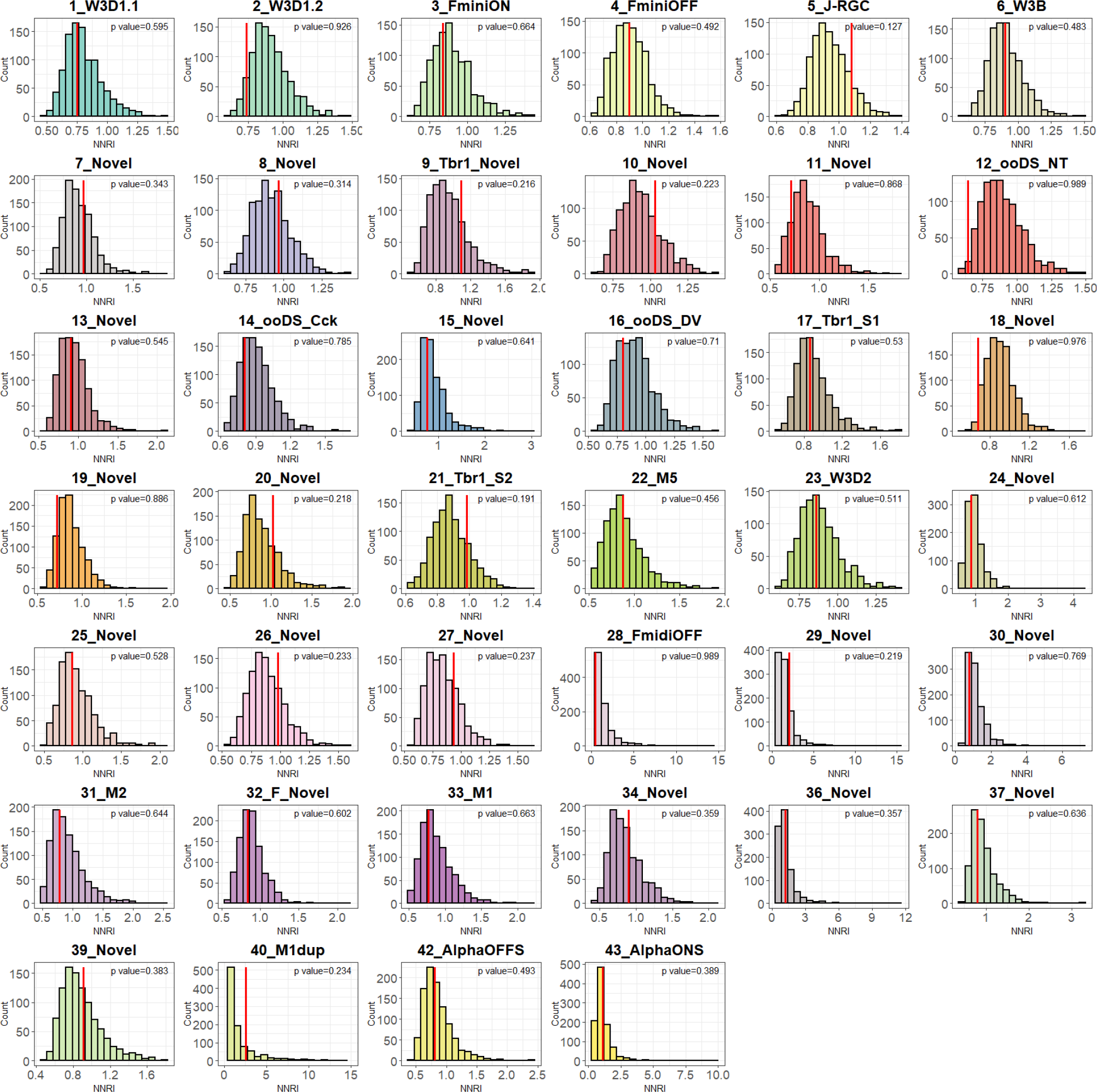
Comparison between observed and random regularity indexes of RGC subtypes. Distribution of randomly permuted NNRIs is shown in the histogram and observed NNRI in red lines. All RGC subtypes showed non-significant difference in NNRI (in red line) compared with random permutation.

**Supplemental 8.**
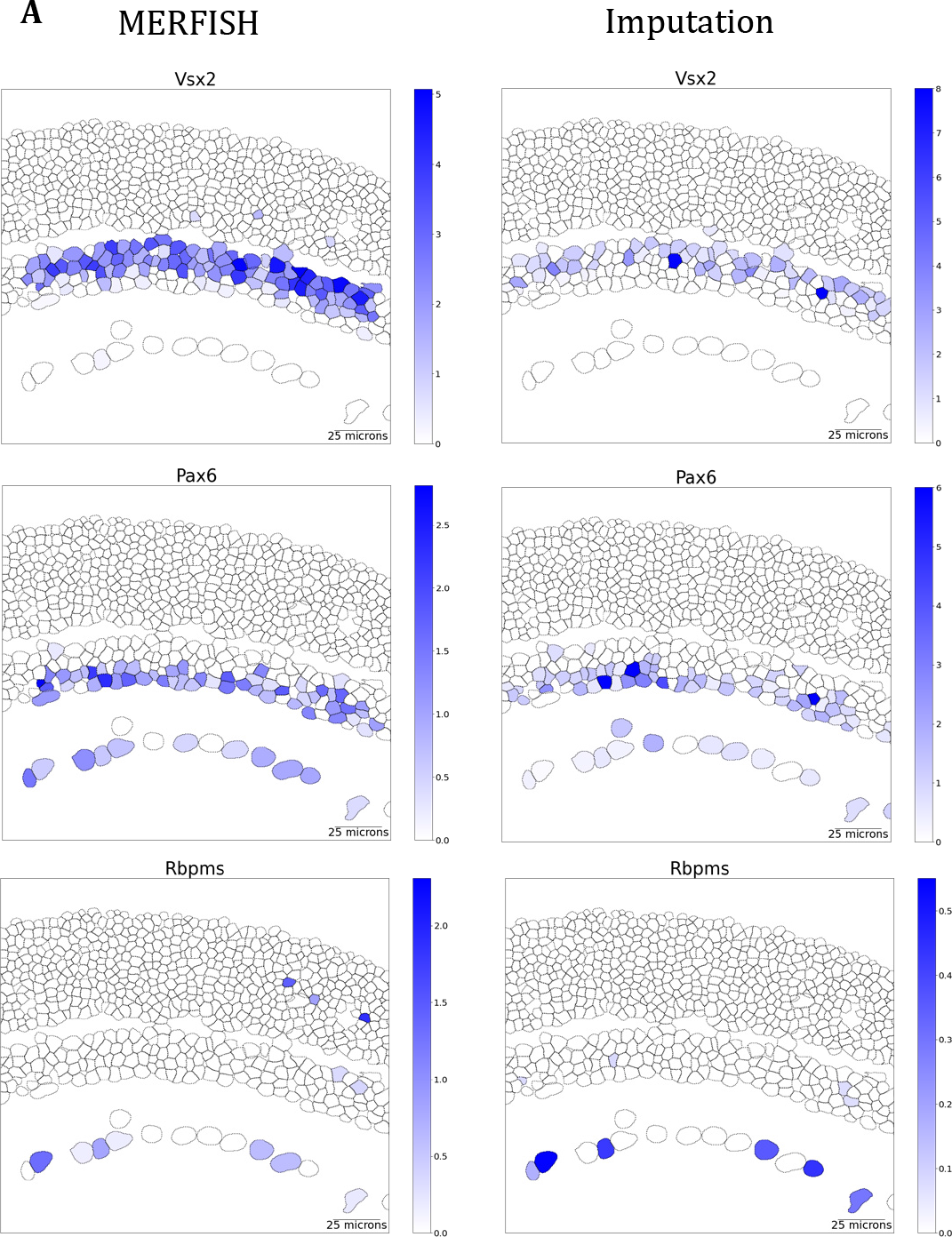
Inferred gene expression, and shared DEGs between three AC subtypes. **(A)** Spatial gene expression patterns of cell type markers in MERFISH profiling and imputation. Cell type markers such as Vsx2 for BC, Pax6 for AC, Rbpms for RGC were profiled in MERFISH experiments, which also show proper spatial pattern in the imputed transcriptome.

**Supplemental 9.**
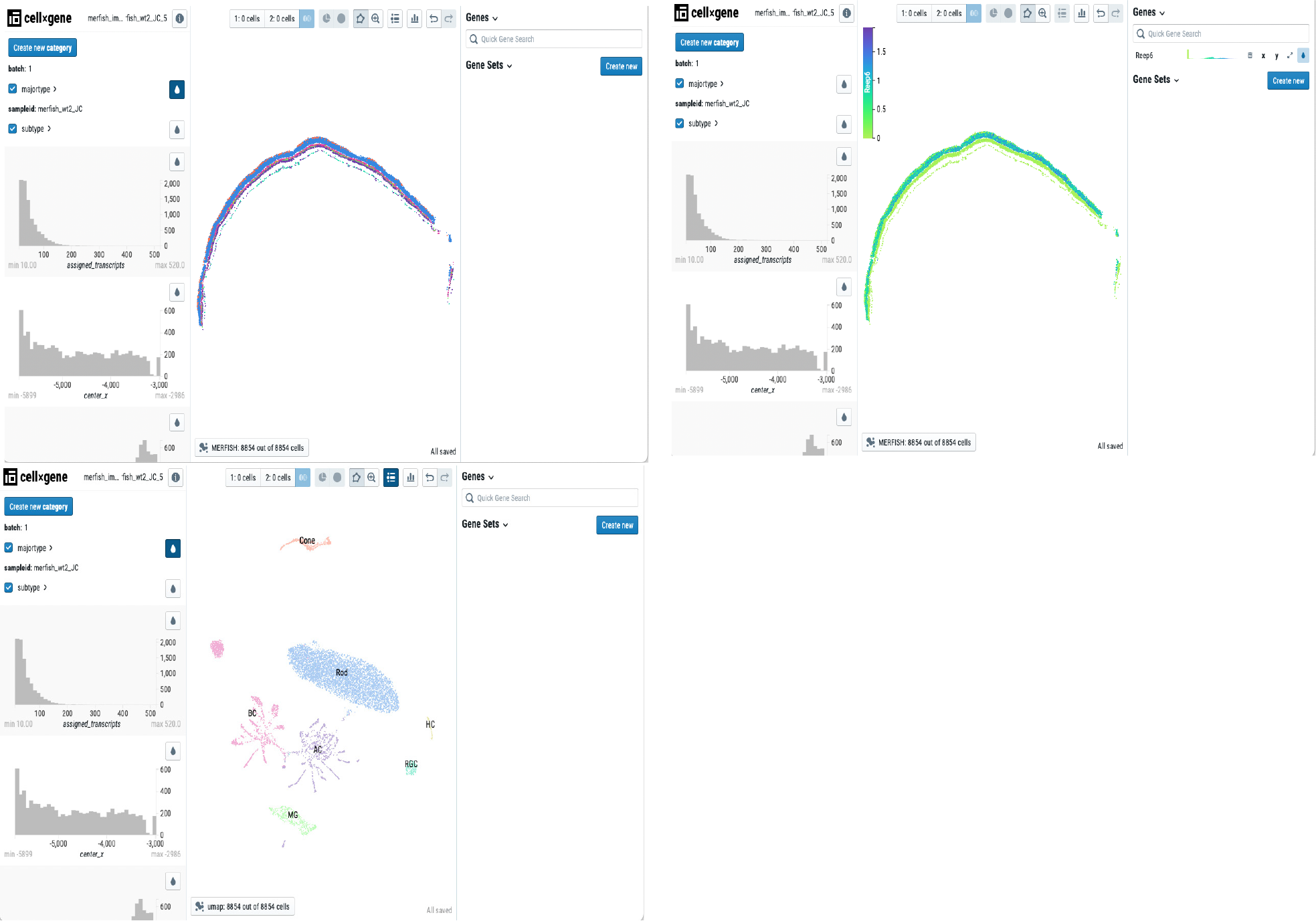
Data resource visualization.

## References

1. Akrouh, A., and Kerschensteiner, D. (2015). Morphology and function of three VIP-expressing amacrine cell types in the mouse retina. J Neurophysiol 114, 2431–2438. https://doi.org/10.1152/jn.00526.2015.

2. Amini, R., Rocha-Martins, M., and Norden, C. (2018). Neuronal Migration and Lamination in the Vertebrate Retina. Front Neurosci 11, 742. https://doi.org/10.3389/fnins.2017.00742.

3. Biancalani, T., Scalia, G., Buffoni, L., Avasthi, R., Lu, Z., Sanger, A., Tokcan, N., Vanderburg, C.R., Segerstolpe, Å., Zhang, M., et al. (2021). Deep learning and alignment of spatially resolved single-cell transcriptomes with Tangram. Nat Methods 18, 1352–1362. https://doi.org/10.1038/s41592-021-01264-7.

4. Bivand, R., Lewin-Koh, N., Pebesma, E., Archer, E., Baddeley, A., Bearman, N., Bibiko, H.-J., Brey, S., Callahan, J., and Carrillo, G. (2022). Package ‘maptools.’

5. Blighe, K., Rana, S., and Lewis, M. (2018). EnhancedVolcano: Publication-ready volcano plots with enhanced colouring and labeling.

6. Bradski, G. (2000). The OpenCV Library. Dr. Dobb’s Journal of Software Tools.

7. Burger, C.A., Albrecht, N.E., Jiang, D., Liang, J.H., Poché, R.A., and Samuel, M.A. (2021). LKB1 and AMPK instruct cone nuclear position to modify visual function. Cell Reports 34, 108698. https://doi.org/10.1016/j.celrep.2021.108698.

8. Casella, G., and Berger, R.L. (2021). Statistical inference (Cengage Learning).

9. Chang, W., Cheng, J., Allaire, J.J., Sievert, C., Schloerke, B., Xie, Y., Allen, J., McPherson, J., Dipert, A., and Borges, B. (2021). shiny: Web Application Framework for R.

10. Chen, K.H., Boettiger, A.N., Moffitt, J.R., Wang, S., and Zhuang, X. (2015). Spatially resolved, highly multiplexed RNA profiling in single cells. Science 348, aaa6090. https://doi.org/10.1126/science.aaa6090.

11. Chen, Y., Song, J., Ruan, Q., Zeng, X., Wu, L., Cai, L., Wang, X., and Yang, C. (2021). Single-Cell Sequencing Methodologies: From Transcriptome to Multi-Dimensional Measurement. Small Methods 5, 2100111. https://doi.org/10.1002/smtd.202100111.

12. Cherry, T.J., Trimarchi, J.M., Stadler, M.B., and Cepko, C.L. (2009). Development and diversification of retinal amacrine interneurons at single cell resolution. Proc Natl Acad Sci U S A 106, 9495–9500. https://doi.org/10.1073/pnas.0903264106.

13. Cho, C.-S., Xi, J., Si, Y., Park, S.-R., Hsu, J.-E., Kim, M., Jun, G., Kang, H.M., and Lee, J.H. (2021). Microscopic examination of spatial transcriptome using Seq-Scope. Cell 184, 3559–3572.e22. https://doi.org/10.1016/j.cell.2021.05.010.

14. Clark, B.S., Stein-O’Brien, G.L., Shiau, F., Cannon, G.H., Davis-Marcisak, E., Sherman, T., Santiago, C.P., Hoang, T.V., Rajaii, F., James-Esposito, R.E., et al. (2019). Single-Cell RNA-Seq Analysis of Retinal Development Identifies NFI Factors as Regulating Mitotic Exit and Late-Born Cell Specification. Neuron 102, 1111–1126.e5. https://doi.org/10.1016/j.neuron.2019.04.010.

15. Della Santina, L., Kuo, S.P., Yoshimatsu, T., Okawa, H., Suzuki, S.C., Hoon, M., Tsuboyama, K., Rieke, F., and Wong, R.O.L. (2016). Glutamatergic Monopolar Interneurons Provide a Novel Pathway of Excitation in the Mouse Retina. Curr Biol 26, 2070–2077. https://doi.org/10.1016/j.cub.2016.06.016.

16. Dou, J., Liang, S., Mohanty, V., Miao, Q., Huang, Y., Liang, Q., Cheng, X., Kim, S., Choi, J., Li, Y., et al. (2022). Bi-order multimodal integration of single-cell data. Genome Biology 23, 112. https://doi.org/10.1186/s13059-022-02679-x.

17. Dries, R., Chen, J., Rossi, N. del, Khan, M.M., Sistig, A., and Yuan, G.-C. (2021a). Advances in spatial transcriptomic data analysis. Genome Res. 31, 1706–1718. https://doi.org/10.1101/gr.275224.121.

18. Dries, R., Zhu, Q., Dong, R., Eng, C.-H.L., Li, H., Liu, K., Fu, Y., Zhao, T., Sarkar, A., Bao, F., et al. (2021b). Giotto: a toolbox for integrative analysis and visualization of spatial expression data. Genome Biology 22, 78. https://doi.org/10.1186/s13059-021-02286-2.

19. Emanuel, G., seichhorn, Babcock, H., leonardosepulveda, and timblosser (2020). ZhuangLab/MERlin: MERlin v0.1.6 (Zenodo).

20. Famiglietti, E.V. (1983). On and off pathways through amacrine cells in mammalian retina: the synaptic connections of “starburst” amacrine cells. Vision Res 23, 1265–1279. https://doi.org/10.1016/0042-6989(83)90102-5.

21. Galli-Resta, L., Novelli, E., Volpini, M., and Strettoi, E. (2000). The spatial organization of cholinergic mosaics in the adult mouse retina. Eur J Neurosci 12, 3819–3822. https://doi.org/10.1046/j.1460-9568.2000.00280.x.

22. Greenwald, N.F., Miller, G., Moen, E., Kong, A., Kagel, A., Dougherty, T., Fullaway, C.C., McIntosh, B.J., Leow, K.X., Schwartz, M.S., et al. (2022). Whole-cell segmentation of tissue images with human-level performance using large-scale data annotation and deep learning. Nat Biotechnol 40, 555–565. https://doi.org/10.1038/s41587-021-01094-0.

23. Greferath, U., Grünert, U., and Wässle, H. (1990). Rod bipolar cells in the mammalian retina show protein kinase C-like immunoreactivity. J Comp Neurol 301, 433–442. https://doi.org/10.1002/cne.903010308.

24. Haverkamp, S., Ghosh, K.K., Hirano, A.A., and Wässle, H. (2003). Immunocytochemical description of five bipolar cell types of the mouse retina. J Comp Neurol 455, 463–476. https://doi.org/10.1002/cne.10491.

25. Jacoby, J., Zhu, Y., DeVries, S.H., and Schwartz, G.W. (2015). An Amacrine Cell Circuit for Signaling Steady Illumination in the Retina. Cell Reports 13, 2663–2670. https://doi.org/10.1016/j.celrep.2015.11.062.

26. Jeon, C.J., Strettoi, E., and Masland, R.H. (1998). The major cell populations of the mouse retina. J Neurosci 18, 8936–8946. .

27. Jeon, Y.-K., Kim, T.-J., Lee, J.-Y., Choi, J.-S., and Jeon, C.-J. (2007). AII amacrine cells in the inner nuclear layer of bat retina: identification by parvalbumin immunoreactivity. Neuroreport 18, 1095–1099. https://doi.org/10.1097/WNR.0b013e3281e72afe.

28. Kay, J.N., Voinescu, P.E., Chu, M.W., and Sanes, J.R. (2011). Neurod6 expression defines new retinal amacrine cell subtypes and regulates their fate. Nat Neurosci 14, 965–972. https://doi.org/10.1038/nn.2859.

29. Kay, J.N., Chu, M.W., and Sanes, J.R. (2012). MEGF10 and MEGF11 mediate homotypic interactions required for mosaic spacing of retinal neurons. Nature 483, 465–469. https://doi.org/10.1038/nature10877.

30. Keeley, P.W., Eglen, S.J., and Reese, B.E. (2020). From random to regular: Variation in the patterning of retinal mosaics*. Journal of Comparative Neurology 528, 2135–2160. https://doi.org/10.1002/cne.24880.

31. Lopez, R., Regier, J., Cole, M.B., Jordan, M.I., and Yosef, N. (2018). Deep generative modeling for single-cell transcriptomics. Nat Methods 15, 1053–1058. https://doi.org/10.1038/s41592-018-0229-2.

32. Love, M.I., Huber, W., and Anders, S. (2014). Moderated estimation of fold change and dispersion for RNA-seq data with DESeq2. Genome Biology 15, 550. https://doi.org/10.1186/s13059-014-0550-8.

33. Lu, Y., Liu, M., Yang, J., Weissman, S.M., Pan, X., Katz, S.G., and Wang, S. (2021). Spatial transcriptome profiling by MERFISH reveals fetal liver hematopoietic stem cell niche architecture. Cell Discov 7, 1–17. https://doi.org/10.1038/s41421-021-00266-1.

34. Macosko, E.Z., Basu, A., Satija, R., Nemesh, J., Shekhar, K., Goldman, M., Tirosh, I., Bialas, A.R., Kamitaki, N., Martersteck, E.M., et al. (2015). Highly parallel genome-wide expression profiling of individual cells using nanoliter droplets. Cell 161, 1202–1214. https://doi.org/10.1016/j.cell.2015.05.002.

35. McInnes, L., Healy, J., Saul, N., and Großberger, L. (2018). UMAP: Uniform Manifold Approximation and Projection. Journal of Open Source Software 3, 861. https://doi.org/10.21105/joss.00861.

36. Megill, C., Martin, B., Weaver, C., Bell, S., Prins, L., Badajoz, S., McCandless, B., Pisco, A.O., Kinsella, M., Griffin, F., et al. (2021). cellxgene: a performant, scalable exploration platform for high dimensional sparse matrices. 2021.04.05.438318. https://doi.org/10.1101/2021.04.05.438318.

37. Moffitt, J.R., Bambah-Mukku, D., Eichhorn, S.W., Vaughn, E., Shekhar, K., Perez, J.D., Rubinstein, N.D., Hao, J., Regev, A., Dulac, C., et al. (2018). Molecular, Spatial and Functional Single-Cell Profiling of the Hypothalamic Preoptic Region. Science 362, eaau5324. https://doi.org/10.1126/science.aau5324.

38. Morrow, E.M., Chen, C.-M.A., and Cepko, C.L. (2008). Temporal order of bipolar cell genesis in the neural retina. Neural Development 3, 2. https://doi.org/10.1186/1749-8104-3-2.

39. Nemitz, L., Dedek, K., and Janssen-Bienhold, U. (2021). Synaptic Remodeling in the Cone Pathway After Early Postnatal Horizontal Cell Ablation. Front Cell Neurosci 15, 657594. https://doi.org/10.3389/fncel.2021.657594.

40. Pang, J.-J., and Wu, S.M. (2011). Morphology and Immunoreactivity of Retrogradely Double-Labeled Ganglion Cells in the Mouse Retina. Investigative Ophthalmology & Visual Science 52, 4886–4896. https://doi.org/10.1167/iovs.10-5921.

41. Pateiro-López, B., and Casal, A. (2010). Generalizing the Convex Hull of a Sample: The R Package alphahull. Journal of Statistical Software 34, 1–28. https://doi.org/10.18637/jss.v034.i05.

42. Peng, Y.-R., James, R.E., Yan, W., Kay, J.N., Kolodkin, A.L., and Sanes, J.R. (2020). Binary Fate Choice between Closely Related Interneuronal Types Is Determined by a Fezf1-Dependent Postmitotic Transcriptional Switch. Neuron 105, 464–474.e6. https://doi.org/10.1016/j.neuron.2019.11.002.

43. Rao, A., Barkley, D., França, G.S., and Yanai, I. (2021). Exploring tissue architecture using spatial transcriptomics. Nature 596, 211–220. https://doi.org/10.1038/s41586-021-03634-9.

44. Raven, M.A., and Reese, B.E. (2002). Horizontal cell density and mosaic regularity in pigmented and albino mouse retina. J Comp Neurol 454, 168–176. https://doi.org/10.1002/cne.10444.

45. Reese, B.E., and Keeley, P.W. (2015). Design principles and developmental mechanisms underlying retinal mosaics. Biol Rev Camb Philos Soc 90, 854–876. https://doi.org/10.1111/brv.12139.

46. Reese, B.E., and Keeley, P.W. (2016). Genomic Control of Neuronal Demographics in the Retina. Prog Retin Eye Res 55, 246–259. https://doi.org/10.1016/j.preteyeres.2016.07.003.

47. Shekhar, K., Lapan, S.W., Whitney, I.E., Tran, N.M., Macosko, E.Z., Kowalczyk, M., Adiconis, X., Levin, J.Z., Nemesh, J., Goldman, M., et al. (2016). Comprehensive Classification of Retinal Bipolar Neurons by Single-Cell Transcriptomics. Cell 166, 1308–1323.e30. https://doi.org/10.1016/j.cell.2016.07.054.

48. Sonntag, S., Dedek, K., Dorgau, B., Schultz, K., Schmidt, K.-F., Cimiotti, K., Weiler, R., Löwel, S., Willecke, K., and Janssen-Bienhold, U. (2012). Ablation of Retinal Horizontal Cells from Adult Mice Leads to Rod Degeneration and Remodeling in the Outer Retina. J. Neurosci. 32, 10713–10724. https://doi.org/10.1523/JNEUROSCI.0442-12.2012.

49. Stringer, C., Wang, T., Michaelos, M., and Pachitariu, M. (2021). Cellpose: a generalist algorithm for cellular segmentation. Nat Methods 18, 100–106. https://doi.org/10.1038/s41592-020-01018-x.

50. Tauchi, M., and Masland, R. (1985). Local order among the dendrites of an amacrine cell population. J Neurosci 5, 2494–2501. https://doi.org/10.1523/JNEUROSCI.05-09-02494.1985.

51. Traag, V.A., Waltman, L., and van Eck, N.J. (2019). From Louvain to Leiden: guaranteeing well-connected communities. Sci Rep 9, 5233. https://doi.org/10.1038/s41598-019-41695-z.

52. Tran, N.M., Shekhar, K., Whitney, I.E., Jacobi, A., Benhar, I., Hong, G., Yan, W., Adiconis, X., Arnold, M.E., Lee, J.M., et al. (2019). Single-Cell Profiles of Retinal Ganglion Cells Differing in Resilience to Injury Reveal Neuroprotective Genes. Neuron 104, 1039–1055.e12. https://doi.org/10.1016/j.neuron.2019.11.006.

53. Vaney, D.I. (1990). Chapter 2 The mosaic of amacrine cells in the mammalian retina. Progress in Retinal Research 9, 49–100. https://doi.org/10.1016/0278-4327(90)90004-2.

54. Vaney, D.I., Sivyer, B., and Taylor, W.R. (2012). Direction selectivity in the retina: symmetry and asymmetry in structure and function. Nat Rev Neurosci 13, 194–208. https://doi.org/10.1038/nrn3165.

55. Virtanen, P., Gommers, R., Oliphant, T.E., Haberland, M., Reddy, T., Cournapeau, D., Burovski, E., Peterson, P., Weckesser, W., Bright, J., et al. (2020). SciPy 1.0: fundamental algorithms for scientific computing in Python. Nat Methods 17, 261–272. https://doi.org/10.1038/s41592-019-0686-2.

56. Voinescu, P.E., Emanuela, P., Kay, J.N., and Sanes, J.R. (2009). Birthdays of retinal amacrine cell subtypes are systematically related to their molecular identity and soma position. J Comp Neurol 517, 737–750. https://doi.org/10.1002/cne.22200.

57. Wang, F., Flanagan, J., Su, N., Wang, L.-C., Bui, S., Nielson, A., Wu, X., Vo, H.-T., Ma, X.-J., and Luo, Y. (2012). RNAscope. J Mol Diagn 14, 22–29. https://doi.org/10.1016/j.jmoldx.2011.08.002.

58. Wässle, H., Riemann, H.J., and Boycott, B.B. (1978). The mosaic of nerve cells in the mammalian retina. Proceedings of the Royal Society of London. Series B. Biological Sciences 200, 441–461. https://doi.org/10.1098/rspb.1978.0026.

59. West, E.R., and Cepko, C.L. (2022). Development and diversification of bipolar interneurons in the mammalian retina. Developmental Biology 481, 30–42. https://doi.org/10.1016/j.ydbio.2021.09.005.

60. West, E.R., Lapan, S.W., Lee, C., Kajderowicz, K.M., Li, X., and Cepko, C.L. (2022). Spatiotemporal patterns of neuronal subtype genesis suggest hierarchical development of retinal diversity. Cell Reports 38, 110191. https://doi.org/10.1016/j.celrep.2021.110191.

61. Whitney, I.E., Keeley, P.W., Raven, M.A., and Reese, B.E. (2008). Spatial patterning of cholinergic amacrine cells in the mouse retina. J Comp Neurol 508, 1–12. https://doi.org/10.1002/cne.21630.

62. Wolf, F.A., Angerer, P., and Theis, F.J. (2018). SCANPY: large-scale single-cell gene expression data analysis. Genome Biology 19, 15. https://doi.org/10.1186/s13059-017-1382-0.

63. Yan, W., Laboulaye, M.A., Tran, N.M., Whitney, I.E., Benhar, I., and Sanes, J.R. (2020). Mouse Retinal Cell Atlas: Molecular Identification of over Sixty Amacrine Cell Types. J Neurosci 40, 5177–5195. https://doi.org/10.1523/JNEUROSCI.0471-20.2020.

64. Zhang, M., Eichhorn, S.W., Zingg, B., Yao, Z., Cotter, K., Zeng, H., Dong, H., and Zhuang, X. (2021). Spatially resolved cell atlas of the mouse primary motor cortex by MERFISH. Nature 598, 137–143. https://doi.org/10.1038/s41586-021-03705-x.

65. Zhu, Y., Xu, J., Hauswirth, W.W., and DeVries, S.H. (2014). Genetically Targeted Binary Labeling of Retinal Neurons. J Neurosci 34, 7845–7861. https://doi.org/10.1523/JNEUROSCI.2960-13.2014.

66. Zhuang, X. (2021). Spatially resolved single-cell genomics and transcriptomics by imaging. Nat Methods 18, 18–22. https://doi.org/10.1038/s41592-020-01037-8.

